# Extensive T1-weighted MRI Preprocessing Improves Generalizability of Deep Brain Age Prediction Models^⋆^

**DOI:** 10.1101/2023.05.10.540134

**Authors:** Lara Dular, Franjo Pernuš, Žiga Špiclin

**Author notes:** This document is the results of the research project funded by the Slovenian Research Agency (Core Research Grant No. P2-0232 and Research Grants Nos. J2-2500 and J2-3059). (L. Dular); (Ž. Špiclin).

## Abstract

Brain age is an estimate of chronological age obtained from T1-weighted magnetic resonance images (T1w MRI) and represents a simple diagnostic biomarker of brain ageing and associated diseases. While the current best accuracy of brain age predictions on T1w MRIs of healthy subjects ranges from two to three years, comparing results from different studies is challenging due to differences in the datasets, T1w preprocessing pipelines, and performance metrics used. This paper investigates the impact of T1w image preprocessing on the performance of four deep learning brain age models presented in recent literature. Four preprocessing pipelines were evaluated, differing in terms of registration, grayscale correction, and software implementation. The results showed that the choice of software or preprocessing steps can significantly affect the prediction error, with a maximum increase of 0.7 years in mean absolute error (MAE) for the same model and dataset. While grayscale correction had no significant impact on MAE, the affine registration, compared to the rigid registration of T1w images to brain atlas was shown to statistically significantly improve MAE. Models trained on 3D images with isotropic 1 *mm*^3^ resolution exhibited less sensitivity to the T1w preprocessing variations compared to 2D models or those trained on downsampled 3D images. Some proved invariant to the preprocessing pipeline, however only after offset correction. Our findings generally indicate that extensive T1w preprocessing enhances the MAE, especially when applied to a new dataset. This runs counter to prevailing research literature which suggests that models trained on minimally preprocessed T1w scans are better poised for age predictions on MRIs from unseen scanners. Regardless of model or T1w preprocessing used, we show that to enable generalization of model’s performance on a new dataset with either the same or different T1w preprocessing than the one applied in model training, some form of offset correction should be applied.

**Highlights:**

- Intensive preprocessing improves performance of computationally less demanding models.
- Models trained on 1*mm*^3^ MRIs are most insensitive to the choice of preprocessing.
- Change in preprocessing increased MAE up to 0.7 years for the same model and dataset.
- Preprocessing software used may impact model performance.
- Prediction bias is systematic across the whole age span and inherent to each model.

## 1. Introduction

In the last decade, brain age has emerged as an pivotal measure of biological age, aiming to uncover patterns and heterogeneity of the ageing process. As a neurological biomarker of individual brain health (Franke and Gaser, 2019), brain age may deviate from an individual’s chronological age even for healthy individuals. Multiple research has shown an association of increased brain age with various environmental and lifestyle factors, such as tobacco and alcohol consumption (Ning et al., 2020; Linli et al., 2022; Cole, 2020; Franke et al., 2013), where as physical activity was shown to reduce brain age (Bittner et al., 2021). Furthermore, brain age has been associated with various factors including cognition (Cole, 2020; Smith et al., 2019; Habes et al., 2021; Jawinski et al., 2022), lower verbal fluency scores (Franke et al., 2013), cognitive impairment (Liem et al., 2017) and even social isolation (Lay-Yee et al., 2023).

In the presence of a certain pathology or chronic disease the difference or gap between brain age and chronological age is likely further increased. For instance, premature brain ageing has been demonstrated in neurological diseases and disorders like the Alzheimer’s dementia (Franke and Gaser, 2012), multiple sclerosis (Høgestøl et al., 2019; Cole et al., 2020) schizophrenia (Schnack et al., 2016; Koutsouleris et al., 2014), and other diseases, such as type 2 diabetes (Franke et al., 2013), infection with human immunodeficiency virus (HIV) (Petersen et al., 2021; Cole et al., 2017), and in young adults after premature birth (Hedderich et al., 2021). Evaluating the age gap thus represents an evolving diagnostic biomarker, opening an avenue for researchers to disentangle patterns of brain ageing and associated diseases.

While primary efforts in this domain rely on regression models to predict age from T1-weighted (T1w) magnetic resonance imaging (MRI), the field’s scope has broadened to include other neuroimaging modalities, such as diffusion tensor imaging (Richard et al., 2018), T2 (Liem et al., 2017; Cole, 2020; Hwang et al., 2021), T2-FLAIR (Cole, 2020), as well as functional MRI (Liem et al., 2017; Cole, 2020; Gao et al., 2023).

Since the inception of brain age research, the initial focus primarily revolved around utilizing traditional machine learning models. Though traditional machine learning models maintain a foothold in the field, there has been a discernible shift towards the adoption of deep learning models (Baecker et al., 2021), due to the increasing number of publicly available T1w MRI datasets, large private datasets and the overall popularity and success of deep learning algorithms. As opposed to standard machine learning methods, deep learning allows us to train models directly on MRIs using minimal preprocessing.

Increasingly more accurate age predictions were achieved using various combinations of convolutional neural network (CNN) model architectures, image preprocessing, training strategies, etc. However, due to differences in MRI preprocessing pipelines and software implementations used, it is difficult to disentangle the contribution of methodological innovations from the impact of preprocessing. Furthermore, there is a lack of rigorous statistical analysis to consider the many confounding factors such as the level of grayscale corrections applied, the number of degrees of freedom in T1w to atlas co-registration, software implementation, model architecture and subject/dataset variability, to name a few, rendering an objective comparison between different brain age studies rather difficult.

This paper is organized as follows: a review of related work and our contributions are given in Section 2. In Section 3 we describe the datasets, preprocesing procedures, and deep learning models for brain age prediction. The evaluation protocol is described in Section 4.1, the experiments with results in Sections 4.2, 4.3 and 4.4. Finally, discussion and conclusion are given in respective Sections 5 and 6.

## 2. Related Work and Our Contributions

We focus our background review on brain age prediction literature involving the use of deep learning models. Generally, these are Convolutional Neural Networks (CNN) that input T1w MRIs and are trained to output a scalar value or interval corresponding to the subject’s age. Previous studies differ substantially in the number of subject scans, their age span and the nature and level of applied image preprocessing.

Preprocessing pipelines used in brain age studies generally include gray scale enhancement, such as bias field corrections (Lam et al., 2020; Peng et al., 2021; Dufumier et al., 2021; Feng et al., 2020), and registration to a brain atlas. Registration’s degrees of freedom varied, with studies using rigid (Dartora et al., 2023; Cole et al., 2017), linear (Lam et al., 2020; Peng et al., 2021; Dufumier et al., 2021; Ueda et al., 2019; Huang et al., 2017; Feng et al., 2020) or even nonlinear transforms (Bintsi et al., 2020; Peng et al., 2021; Cheng et al., 2021). Skull stripping that involves extracting the brain from surrounding tissues was also applied in certain studies (Bintsi et al., 2020; Lam et al., 2020; Fisch et al., 2021; Dufumier et al., 2021; Feng et al., 2020; Cheng et al., 2021). In the presence of such preprocessing variations it is difficult to compare study results and disentangle the factors contributing to the accuracy of brain age predictions. A comprehensive study on natural images found that the effect of image preprocessing and augmentation on prediction model performance was greater than the effects of variability in prediction model architecture (Lathuilière et al., 2020), which highlights the need for further research in this area in order to standardize the T1w preprocessing methods for best accuracy of brain age prediction models.

Besides training on the T1w MRIs, brain age models are often trained on segmentation of Gray Matter (GM) and White Matter (WM) structures. Cole et al. (2017) compared models trained on GM segmentations with the model trained on T1w MRI without grayscale corrections, which was rigidly registered to Montreal Neurological Institute (MNI) 152 brain atlas. They found that models trained on GM, with mean absolute error (MAE) of 4.16 years, performed better than models trained on T1w with MAE 4.65 and WM images with MAE 5.14. Better performance on GM than on WM segmentations was confirmed by Peng et al. (2021). They further compared models trained on T1w MRIs with bias field correction with both linear and non-linear spatial registration to the MNI brain atlas. The non-linear registration achieved lower MAE of 2.73 years, which was comparable to accuracy achieved by models trained with GM segmentations for linearly registered T1w image with MAE equal to 2.80 years. Finally, Dufumier et al. (2021) reported comparable results of brain age models based on T1w and GM segmentations when testing on same site images, however, the results reported on an independent new-site test set, not used during model training, show preference to models based on GM segmentations.

Differences in T1w preprocessing may arise from the use of different software implementations (Fisch et al., 2021). Related neuroimaging studies show that the measurements of cortical surface thickness differ significantly between pipelines (Kharabian Masouleh et al., 2020; Bhagwat et al., 2021) and reveal a significant discrepancy between the cortical thickness reproducibility metrics (de Fátima Machado Dias et al., 2022). Reasons could also involve T1w MRI resolution variations and contrast-to-noise differences. It seems that the use of GM segmentation for brain age predictions is rather ill-posed and, therefore, this study will focus on preprocessed T1w images as model input. It is yet to be determined if there is a significant effect of the software implementations on brain age prediction even for fairly simple T1w preprocessing operations.

To cut the computational overhead of T1w preprocessing and mitigate potential bias introduced by different software implementations, a recent review paper calls for a further development of brain age from routine MRIs, with minimal preprocessing (Tanveer et al., 2023). The potential and general applicability of such models was already argued by Cole et al. (2017), who proposed one of the first deep learning models for brain age regression. Their model was trained on approximately 2,000 T1w MRIs, with the T1w preprocessing involving only rigid registration to MNI brain atlas, and achieved a MAE of 4.65 years. On a much larger dataset of over 16,000 MRIs using the same minimal T1w preprocessing Dartora et al. (2023) achieved a MAE score of 2.67 years. Further along this line, Fisch et al. (2021) considered minimal T1w preprocessing as applying only skull striping and no spatial registration. Their Residual Network (ResNet) based model, trained on approximately 10,000 MRIs, achieved MAE of 2.84 years. These recent models seem to achieve competitive results in comparison to previously mentioned models, which were trained on datasets of approximately the same size, but with more extensive T1w preprocessing.

Validation of brain age prediction models for clinical application should involve their performance assessment on a dataset from a new (unseen) site, not used during model training. This is a common use case scenario, occurring when applying a pretrained model on new data, possibly preprocessed with a different pipeline. In such scenario, Feng et al. (2020) reported a rather small increase in MAE of 0.15 years, using the same T1w preprocessing on training and test dataset. Multiple other deep learning studies indicate that this increase (or accuracy deterioration) to be much larger. Jonsson et al. (2019) reported an increase in MAE of about 3 and 5 years on two separate new site datasets. Moreover, Dufumier et al. (2021) showed an increase of MAE by at least 2 years for a wide range of CNN architectures, even for CNNs trained on a large dataset with over 10,000 T1w images.

Drop in brain age prediction accuracy was reported also for models trained on datasets with minimal T1w preprocessing. Dartora et al. (2023) reported a 1 and 3 year increase in MAE on two independent datasets. Fisch et al. (2021) report a 5 year increase on three datasets, prior to applying transfer learning. Therein, the CNN model performed worse than three traditional machine learning models based on explicit feature extraction from T1w MRIs. This increase in MAE therefore seems intrinsically connected to the previously unseen dataset and/or new (unseen) preprocessing procedure and not the model’s ability to generalize.

The contributions of this study are the following:

i. A thorough and reproducible quantitative assessment of the impact of four T1w preprocessing variants on brain age prediction accuracy using four recent model architectures.
ii. Rigorous statistical evaluation involving repeated model training with random initialization and use of linear mixed-effects models (LMEMs) encompassing the study of the impact of various confounding factors.
iii. Study of model performance generalization, and strategies for its improvement, on a new site dataset and/or new T1w preprocessing approach and software implementation.

## 3. Datasets and Age Prediction Methodology

### 3.1. Datasets

For studying the effect of image preprocessing on brain age prediction, we created two datasets: the first containing multi-site T1w MRIs for training, validation and testing and the second used solely for testing, which contained new unseen site data. All included subjects were healthy individuals, without previously known neurological diseases, from 18 to 95 years old.

The **multi-site dataset** was gathered from seven publicly available datasets. Most datasets within this collection sourced images from multiple hospitals or sites, utilizing an array of MRI scanners, including those from GE, Siemens, and Philips, operating at 1.5T and 3T field strengths. Exceptions are the OASIS 2 and CamCAN datasets, in which scans were acquired on a single scanner. Due to the integration of these multi-source, multi-site, and multi-vendor datasets, variations in acquisition pipelines are inherently present.

The multi-site dataset included a total of 4428 T1w MRIs of healthy subjects. The gathered images were preprocessed using four different preprocessing pipelines and underwent a visual quality check. Images that did not pass the visual quality check for reasons like motion artifacts, failed preprocessing, etc., were excluded (*N*_*excl*_ = 408). Furthermore, subjects under the age of 18 or with missing age information were discarded (*N*_*disc*_ = 481) and, in case multiple scans per subject were available, a single scan (chronologically the first non-discarded image) was retained. Finally accepted were a total of 2504 T1w MRIs, which were split into train (*N* = 2012), validation (*N* = 245) and test (*N* = 247) datasets. The overall statistics per dataset are given in Supplementary Table 4. For reproducibility reasons the exact subject IDs included in each split are given in Supplementary materials (see Section 5.5).

The **unseen site dataset** was chosen as a subset from the UK Biobank (UKB) dataset. For the purpose of testing, we selected T1w MRI scan of 1493 healthy subjects. All included subjects met the inclusion criteria of not having long-standing illnesses and were required to self-report an overall health rating of *excellent* or *good* at the time of scan acquisition. The dataset was preprocessed using the same four preprocessing pipelines as multi-site dataset. In addition, a fifth preprocessing pipeline, given by the dataset providers, was used to observe the model’s ability to predict not only on previously unseen data, but also on previously unseen preprocessing.

For multi-site and unseen site dataset, the ground truth brain age corresponds to the subject’s chronological age, which was either given by the dataset providers or calculated from the provided date of birth and the MRI acquisition date. For the majority of datasets, including ADNI, CamCAN, CC-359, OASIS 2, and FCON 1000, the age was provided as a rounded figure to the nearest year. The age distribution of the included T1w subject scans per dataset, and the train/validation/test subsets, is provided in Supplementary Materials (Table 4, Figure 7).

### 3.2. Image preprocessing pipelines

We implemented four preprocessing pipelines using a combination of publicly available and in-house software. These pipelines vary on the registration methods, the extent of gray scale corrections, as well as the algorithms and software used. For clarity, a schematic representation of all four pipelines is illustrated in Figure 1.

**Figure 1.**
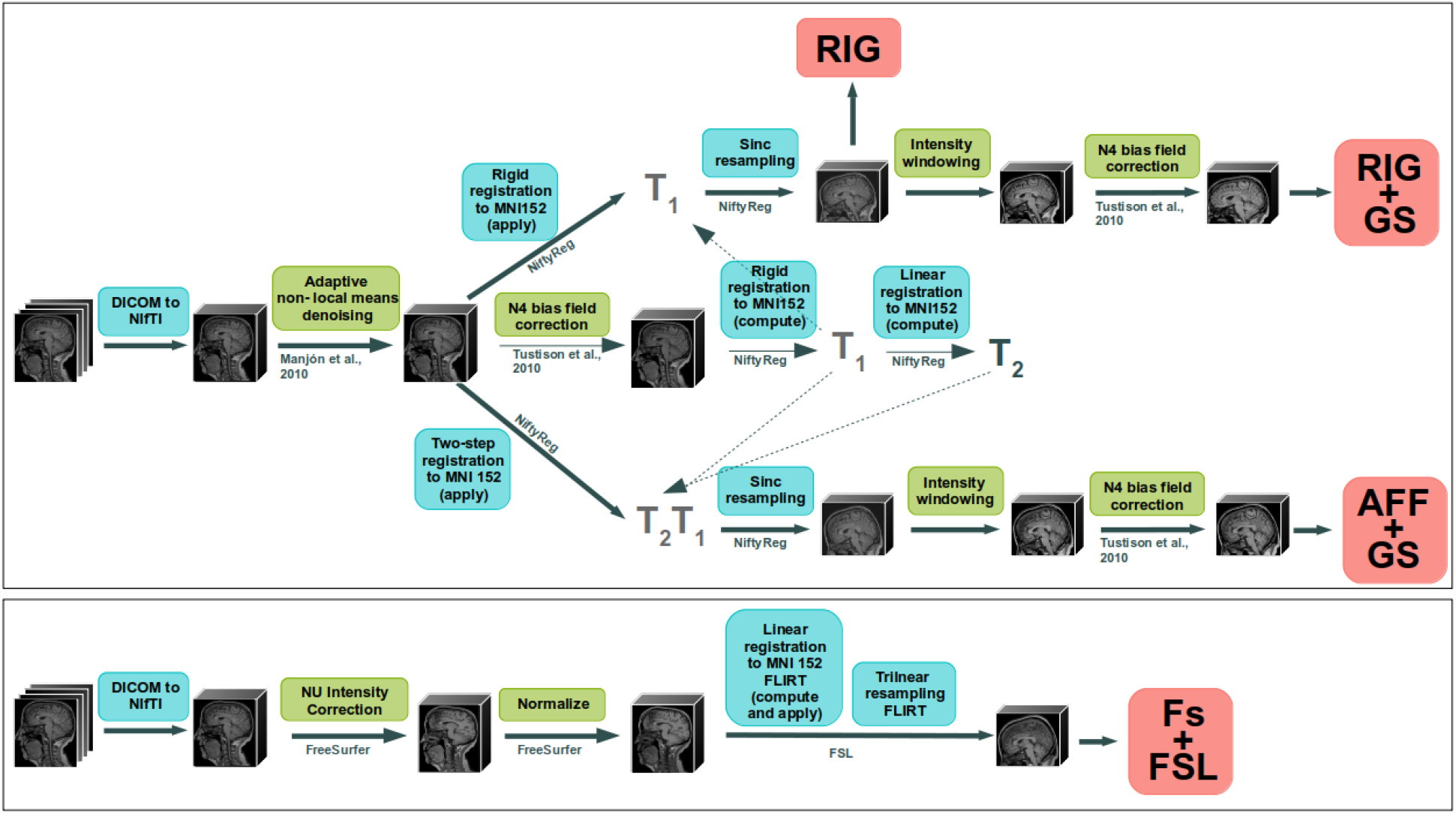
Schematic representation of preprocessing pipelines and software used.

Common to all pipelines, the input T1w image was first converted to the Nifti format. In the first three pipelines, the input raw T1w image was initially denoised using Adaptive non-local means denoising ^1^ with spatially varying noise levels (Manjón et al., 2010).

Aligned with Cole et al. (2017), the first pipeline, denoted **RIG**, performed rigid registration of the denoised T1w image into the MNI152 nonlinear atlas, version 2009c (Fonov et al., 2009), with size 193 × 229 × 193 and spacing 1 *mm*^3^. To improve registration accuracy, intensity inhomogeneity correction (without mask) was applied to the denoised image using N4 algorithm ^2^ (Tustison et al., 2010), prior to running the registration. The inhomogeneity corrected T1w image was used during registration only, while, finally, the denoised T1w image was sinc resampled using the obtained rigid transformation.

The second pipeline, **RIG+GS**, extended the RIG pipeline by applying an additional two-step grayscale correction procedure to the RIG output. The first step, *1)* intensity windowing, involves computation of the lower and upper thresholds based on the grayscale histogram, smoothed with a Gaussian filter. The lower threshold is set based on histogram’s lowest intensity mode location plus twice the value of the mode’s full width at half maximum (FWHM). Note that the particular mode corresponds to the grayscale values of the background and non-tissue regions of the T1w MRI image. To compute the upper threshold, the grayscale values beyond the 99th percentile are first set to the value of the lower threshold. Inflection points in the intensity distribution from the 50th to the 95th percentiles are then identified by computing the second derivative. The upper threshold is defined as the value of the percentile at a selected inflection point, plus three times the Median Absolute Deviation of the pixel intensities that are above the lower threshold. The second step, *2)* involves intensity inhomogeneity correction, utilizing the N4 algorithm with the MNI152 atlas mask dilated by 3 voxels.

The third pipeline, **AFF+GS**, was a modified version of the RIG+GS, by applying in sequence the rigid and affine registration steps. Finally, the two-step grayscale correction procedure was applied as in the RIG+GS pipeline. All previously mentioned image registration and resampling steps for processing pipelines RIG, RIG+GS and AFF+GS were performed using the publicly available NiftyReg software ^3^ (Modat et al., 2014).

The fourth pipeline, **Fs+FSL**, utilized commonly used software tools FreeSurfer^4^ and FSL (FMRIB Software Library) ^5^ (Jenkinson et al., 2012) and included gray scale corrections and affine registration. Raw T1w images were preprocessed using the grayscale correction preprocessing stages of FreeSurfer’s cortical reconstruction `recon-all` pipeline, with default parameter settings. The preprocessing entails non-parametric non-uniform intensity normalization (N3), followed by intensity normalization that sets the mean intensity of the white matter to 110 (Laboratory for Computational Neuroimaging and Athinoula A. Martinos Center for Biomedical Imaging., 2022). In order to ensure consistency among all preprocessing pipelines, we also applied registration to the MNI152 nonlinear atlas, version 2009c (Fonov et al., 2009), the same reference space as used in previous pipelines. Specifically, we used FSL FLIRT (Jenkinson et al., 2002) with default settings, performing linear registration with trilinear resampling.

#### 3.2.1. Adapting UKB preprocessed data

An additional, fifth variant of preprocessed T1w MRIs from the UKB dataset, described in detail by Smith et al. (2020), was obtained from the UKB dataset providers. Namely, from the UKB dataset we included raw T1w defaced images in subject image space, as well as the preprocessed T1w images and corresponding linear transformation matrices that register the raw T1w image to MNI152 nonlinear 6th generation atlas space (Grabner et al., 2006). Since the above four preprocesing procedures involved registration to 7th generation MNI152 atlas, an additional common linear registration between 6th and 7th generation atlas spaces was applied, to assure all images were in the same space.

Each original defaced T1w image was first resampled to the MNI152 nonlinear 6th generation atlas (Grabner et al., 2006) using FSL FLIRT (Jenkinson and Smith, 2001; Jenkinson et al., 2002) and the provided linear transformation matrix, and then linearly registered to the MNI152 nonlinear 7th generation MNI atlas (version 2009, our target space) (Fonov et al., 2009) and resampled using 3rd order interpolation. The linear transformation matrix between the two MNI spaces was pre-computed using FSL’s FLIRT.

### 3.3. Age Prediction Models

To study the effect of preprocessing in relation to model architecture, four fundamentally different CNN models for brain age estimation were reimplemented based on the descriptions in the literature. Only minor alterations, such as adjustments for the input image dimensions, were made to assure comparability across the experiments.

**Model 1**, proposed by Cole et al. (2017), was a convolutional CNN trained on full resolution 3D T1w MRIs. **Model 2**, proposed by Huang et al. (2017), was trained on 2D images by taking 15 equidistantly sampled axial slices as input channels. **Model 3**, proposed by Ueda et al. (2019), was trained on downsampled T1w MRIs. Finally, **Model 4**, proposed by Peng et al. (2021), was a fully convolutional model trained on full resolution 3D images that reported one of best results for brain age prediction among the CNN models. The architectures of the four models are depicted in Figure 2.

**Figure 2.**
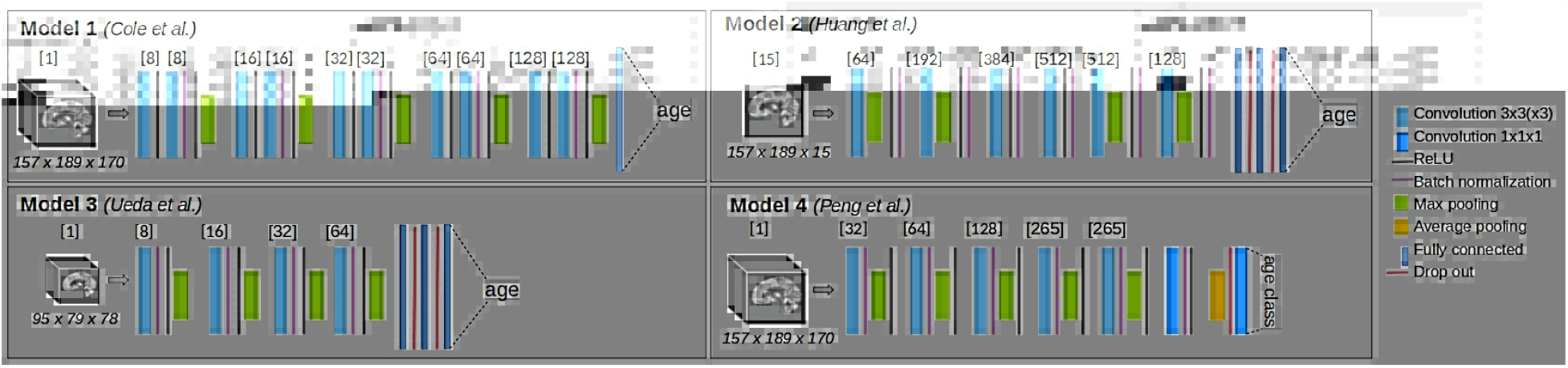
Architecture of the four reimplemented CNN models for brain age prediction.

Brain age estimation is typically formulated as a regression task, such that the model outputs a non-negative real number reflecting the age of the subject based on their T1w MRI scan. Models 1, 2, and 3 therefore had linear activation in the last fully connected layer so as to output the scalar value representing the predicted age.

By contrast, Model 4 was designed as a classification model. Here, the ground truth age value *y* for each sample was ansformed into a so-called *soft label*, represented as Gaussian probability density with mode located at the true age and unit variance. The probability density was discretized into non-overlapping 2-year age intervals by integrating the density over each age interval. The output age prediction was computed as weighted sum over the class probabilities, i.e. *y*′ = Σ_*j*_ *p*_*j*_ *age*_*j*_, where *p*_*j*_ denotes the probability of class *j* and *age*_*j*_ the center of the age class interval.

All models were implemented in PyTorch 1.4.0 for Python 3.6.8.

The model selection was guided by a multifaceted rationale. Firstly, we sought to juxtapose various input MRI representations. By selecting CNN models with distinct input representations, we aimed to assess the influence of preprocessing on models that utilize full-resolution 3D, 2D, and downsampled 3D images. Secondly, by framing age regression as a classification task, as proposed in Model 4, we were intrigued to determine whether such an approach offers inherent stability in the face of preprocessing variations, relative to traditional regression models. Additionally, given the discourse on the computational potential of 2D CNNs in brain age prediction in recent literature (Tanveer et al., 2023) and to reduce computational complexity, our aim was also to ascertain whether models with reduced computational demand, such as downsampled models and 2D models, could attain enhanced performance through extensive image preprocessing.

#### Hyperparameter tuning

The learning rate and batch size hyperparameter values for each model were chosen based on a wide grid search, which was set around the proposed values in corresponding original papers. For instance, tested learning rate values were 10^−2^, 10^−3^, 10^−4^, 5 · 10^−5^, 10^−5^, and 10^−6^. The batch size for Models 2 and 3 was set to 4, 8, 16, 32 and 64. Due to graphics processing unit (GPU) constraints we trained Model 1 with batch size 4, 8, 16 and 24 and Model 4 with batch size 4 and 8. All tested hyperparameter combinations and their results are given in Supplementary Figure 8.

Hyperparameter selection was based on determining the epoch at model convergence, i.e. by observing the course of the loss function, and by observing MAE on the train and validation set in the last 10 epochs. To assure a robust choice of the hyperparameters with respect to both MAE and convergence, we computed median MAE across last 10 training epochs, and the hyperparameter values with smallest median MAE value were chosen as the optimal values.

The chosen optimal hyperparameter values in our study and the originally proposed hyperparameter values are given in Supplementary Table 5. Unless noted otherwise, we used these hyperparameters in all subsequent experiments.

#### Loss function

The choice of loss function depended on the model formulation as either regression or classification network. For Models 1, 2 and 3, we tested both mean squared error (MSE) loss and L1 loss for multiple hyperparameter values. Due to overall better performance and stability of training, all three models were trained with L1 loss. Model 4, defined as a classification model, was trained with Kullback-Leibler divergence loss function.

#### Optimizer

We used the stochastic gradient descent algorithm with momentum 0.9 as proposed in three out of four studies (Cole et al., 2017; Peng et al., 2021), keeping the learning rate decay schedule as originally proposed for each individual model.

We have experimentally determined that Models 1 and 4 typically converged after 110 epochs, while Model 2 and 3 converged after 400 epochs.

#### Data augmentation

All models were trained with the following data augmentation procedures: *1)* random shifting along all major axes with probability of 0.3 for an integer sampled from [−*s, s*], where *s* = 3 for Model 3 (downsampled 3D input T1w) and *s* = 5 for Models 1,2, and 4; *2)* random padding with probability of 0.3 for an integer from range [0, *p*], where *p* = 2 for Model 3 and *p* = 5 for Models 1,2, and 4; *3)* flipping over central sagittal plane with probability of 0.5. Note that the padding and shifting parameters are lower for Model 3, due to the larger image spacing, which is as a result of image downsampling.

Further, the image size as input to the models was adapted during the augmentation. We first we removed the non-informative empty space around the head by cropping to size 157 × 189 × 170 about the image center. Further, for Model 2 the 15 axial slices (predefined in atlas space) were sampled to obtain input image size of 157 × 189 × 15, while for Model 3 the input images were downsampled by a factor of 2 using sinc resampling and cropped to size 95 × 79 × 78.

#### Weighted training

Weighted training is a strategy of assigning higher sampling probabilities to subjects in underrepresented age categories, such that the expected number of samples from each age category becomes equal. Due to the smaller number of subjects in age groups above 80, weighted training was necessary for classification Model 4, but not for the other three models ^6^.

Each subject was assigned a weight of *N*/*n*_*i*_, where *n*_*i*_ denotes the number of samples in category *i*. Age categories were set to [18, 20), [20, 25), [25, 30), …, [85, 90), [90, 100) as previously proposed by Feng et al. (2020) and sampled with replacement. The number of sampled subjects was kept equal to the number of subjects *N*, so that the number per training epoch was kept equal to the experiments without weighted training.

### 3.4. Postprocessing

To enhance prediction accuracy and consistency, we employed model ensembling, averaging multiple age predictions for an individual subject.

Since estimating age on a dataset experiencing domain variation (i.e., new scanner and/or T1w preprocessing) typically results in a decrease in accuracy, manifested as a systematic deviation from the actual age, we applied offset correction when predicting on the UKB dataset. Though bias correction, t. i. fitting a linear regression to predictions on validation or test sets, is commonly used in the literature (Lange et al., 2019; Peng et al., 2021; Cole et al., 2017; Smith et al., 2019; Cheng et al., 2021; Dunås et al., 2021), several recent studies have cautioned against it (Butler et al., 2021; de Lange et al., 2022). Unlike fitting a linear regression line, offset correction does not correct for model’s inability to capture linear trend, nor reduces prediction dispersion.

#### Offset correction

We implemented the offset adjustment by subtracting the value of mean error (ME) from the ensemble prediction, determined as follows:

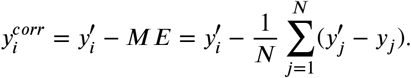

The ME was computed for each model/preprocessing combination. Offset correction was applied only when predicting on new site dataset.

##### Model ensembling

Model ensembling was shown effective in reducing the MAE values, both when combining model outputs obtained from single (Peng et al., 2021; Dufumier et al., 2021; Levakov et al., 2020; Cheng et al., 2021) or multiple preprocesing pipelines (Peng et al., 2021; Kuo et al., 2021).

To avoid reporting the results of a single (possibly lucky) run, each model was trained five times, with different weight initialization. The final prediction of a brain age was obtained as the average of the five model predictions with different weight initialization. On the multi-site T1w train set we trained a total of 80 models: 4× image preprocessing pipelines, 4× model architectures, and 5× random weight initialization.

## 4. Experiments and results

The impact of T1w MRI image preprocessing on the accuracy of brain age predictions using the four CNN models was studies in three scenarios shown in Figure 3: *1)* tested on the same-source dataset and preprocessing as used during model training (**Section 4.2**), *2)* tested on a new unseen dataset but preprocessed in the same way as training dataset, *3)* tested on a new unseen dataset, preprocessed differently than training dataset.

**Figure 3.**
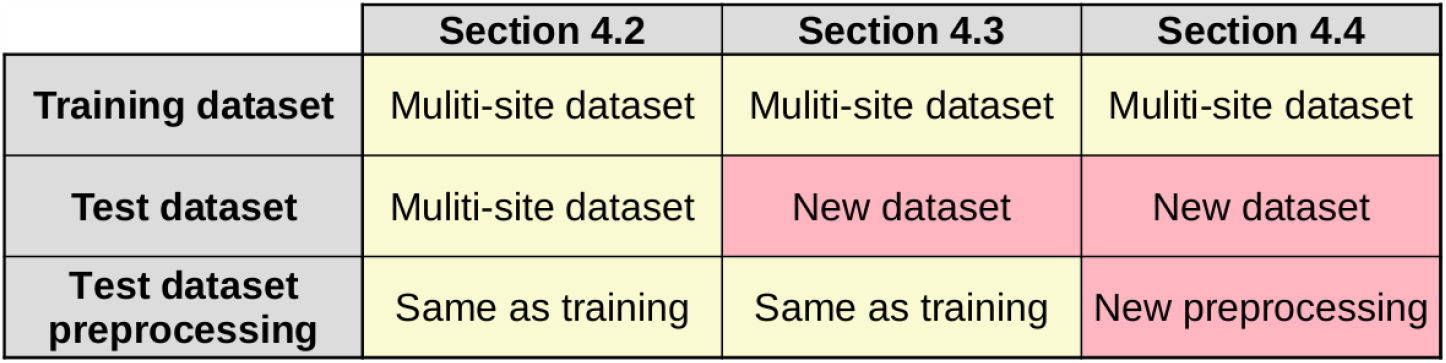
Overview of the tested brain age train and test scenarios.

### 4.1. Evaluation Protocol

For experiment evaluation we computed commonly used performance metrics to highlight specific aspects of the prediction model performances.

Established metric of model accuracy is the mean absolute error (MAE):

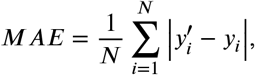

where *y*_*i*_ denotes true age and 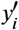 predicted age of *i*-th subject. We also report mean error (ME):

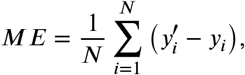

since values of ME deviating from zero show that a model on average either under-or over-estimates age on the whole age interval. Assuming the prediction error is normally distributed around zero, we expect ME to be zero.

#### 4.1.1. Statistical Analysis

LMEMs were used to describe the relationship between a prediction’s absolute error as dependent variable and response variables that were set for each research question. Each LMEM included model architecture, preprocessing procedure and their interaction as fixed effect and subject ID as random effect, such that all responses for a specific subject were shifted by a subject-specific additive value. By modeling subject ID as random effect, we account for dependent data that arises from multiple brain age predictions for the same subject under multiple conditions (preprocessing procedure, model architecture, offset correction).

We employed a stepwise approach in fitting LMEMs. Namely, the models were first constructed with the fixed factors and, subsequently, we incrementally incorporated fixed-factor interactions to increase model complexity. To evaluate the benefit of increasing model complexity, we utilized Analysis of Variance (ANOVA) for model comparison, to test if the increase in complexity resulted in a statistically significant improvement in explaining the observed variability in the data.

For the final LMEM models, we reported regression coefficients and their 95% confidence intervals, provided in Supplementary materials. Results of LMEM analyses were supported by the ANOVA test declaring statistical significance for *p* < 0.01. Further, if the main fixed factor showed a difference in responses, a post-hoc pairwise test was conducted, with confidence level of 0.95, and multiplicity adjustments using Tukey’s correction.

LMEM analysis was conducted in R version 4.0.4, using ‘lme4’ package version 1.1.26. For computing *p*-values of ANOVA tests we used package ‘lmerTest’ version 3.1.3. Finally, pairwise analysis was conducted using package ‘emmeans’ version 1.5.4.

### 4.2. Effect of image preprocessing

Our goal is to evaluate the impact of the particular choice of image preprocessing for various CNN architectures, described in respective Sections 3.2 and 3.3. On the multi-site T1w train set we trained a total of 80 models: 4× image preprocessing pipelines, 4× model architectures, and 5× random weight initializations. Brain age predictions were obtained as the average age prediction of five models trained with different random weight initialization. The model accuracy metrics are presented in Table 1.

**Table 1:** Multi-site test set results for all 16 combinations of preprocessing pipelines and model architectures. Best MAE values wrt. model architecture (*rows*) are marked in **bold**, while best values wrt. image preprocessing procedure (*columns*) are <u>underlined</u>. All numbers are in years.

|  | RIG |  | RIG+GS |  | AFF+GS |  | Fs+FSL |  |
| --- | --- | --- | --- | --- | --- | --- | --- | --- |
|  | ME (sd) | MAE (sd) | ME (sd) | MAE (sd) | ME (sd) | MAE (sd) | ME (sd) | MAE (sd) |
| Model 1 | -0.46 ± 4.10 | <u>3.18 ± 2.61</u> | -0.25 ± 4.14 | <u>3.12 ± 2.73</u> | -0.03 ± 3.86 | **2.96 ± 2.47** | 0.26 ± 4.30 | <u>3.22 ± 2.86</u> |
| Model 2 | -0.96 ± 5.75 | <u>4.47 ± 3.73</u> | -1.11 ± 5.49 | <u>4.29 ± 3.59</u> | -0.80 ± 4.81 | **3.72 ± 3.14** | 0.13 ± 5.44 | <u>4.08 ± 3.58</u> |
| Model 3 | -0.13 ± 4.41 | <u>3.45 ± 2.75</u> | -1.03 ± 4.40 | <u>3.50 ± 2.85</u> | -0.68 ± 3.90 | **3.18 ± 2.35** | -0.26 ± 4.61 | <u>3.45 ± 3.06</u> |
| Model 4 | -0.85 ± 4.33 | <u>3.32 ± 2.90</u> | -0.74 ± 4.34 | <u>3.29 ± 2.92</u> | -0.46 ± 4.21 | **3.25 ± 2.70** | -0.13 ± 4.52 | <u>3.31 ± 3.07</u> |

We further fit a LMEM model with model architecture and preprocessing procedure as main effects and subject ID as random effect. The ANOVA test and 95% CI interval values showed both fixed factors are statistically significant (*F* (3, 3699 = 49.49, *p* < 2.2*e* − 16 for model architecture; *F* (3, 3699) = 5.09, *p* = 0.002 for preprocessing). We increased the LMEM complexity by including the interaction of the fixed factors, however the interaction terms were not statistically significant. Since the main effects were statistically significant (*F* (9, 3699) = 1.14, *p* = 0.328) and the interaction is theoretically meaningful, the interaction was included in the final model despite not being statistically significant. The LMEM coefficients, their 95% CI and ANOVA F-values are reported in Supplementary Table 7.

The results of the LMEM post-hoc pairwise analysis are shown in Figure 4. Model 1 outperformed other models for all preprocessing pipelines (cf. Table 1), however these differences were only significant between Model 1 and Model 2 (cf. Figure 4). Further, the absolute error of Model 2 was found to be significantly higher than MAE of all 3D models (*p* < 0.001), for all but AFF+GS preprocessing. Out of the four, Model 2 has the largest bias, measured by ME, for RIG, RIG+GS and AFF+GS preprocessing pipelines and the largest variability, as evidenced by the higher standard deviation(cf. Table 1). It only achieved MAE below 4 years when trained on the AFF+GS dataset. Notably, it is only with this preprocessing that the performance difference between Model 2 and the other models becomes statistically insignificant for most pairs, as depicted in Figure 4. All of this shows to its poorer performance to the 3D counterparts.

**Figure 4.**
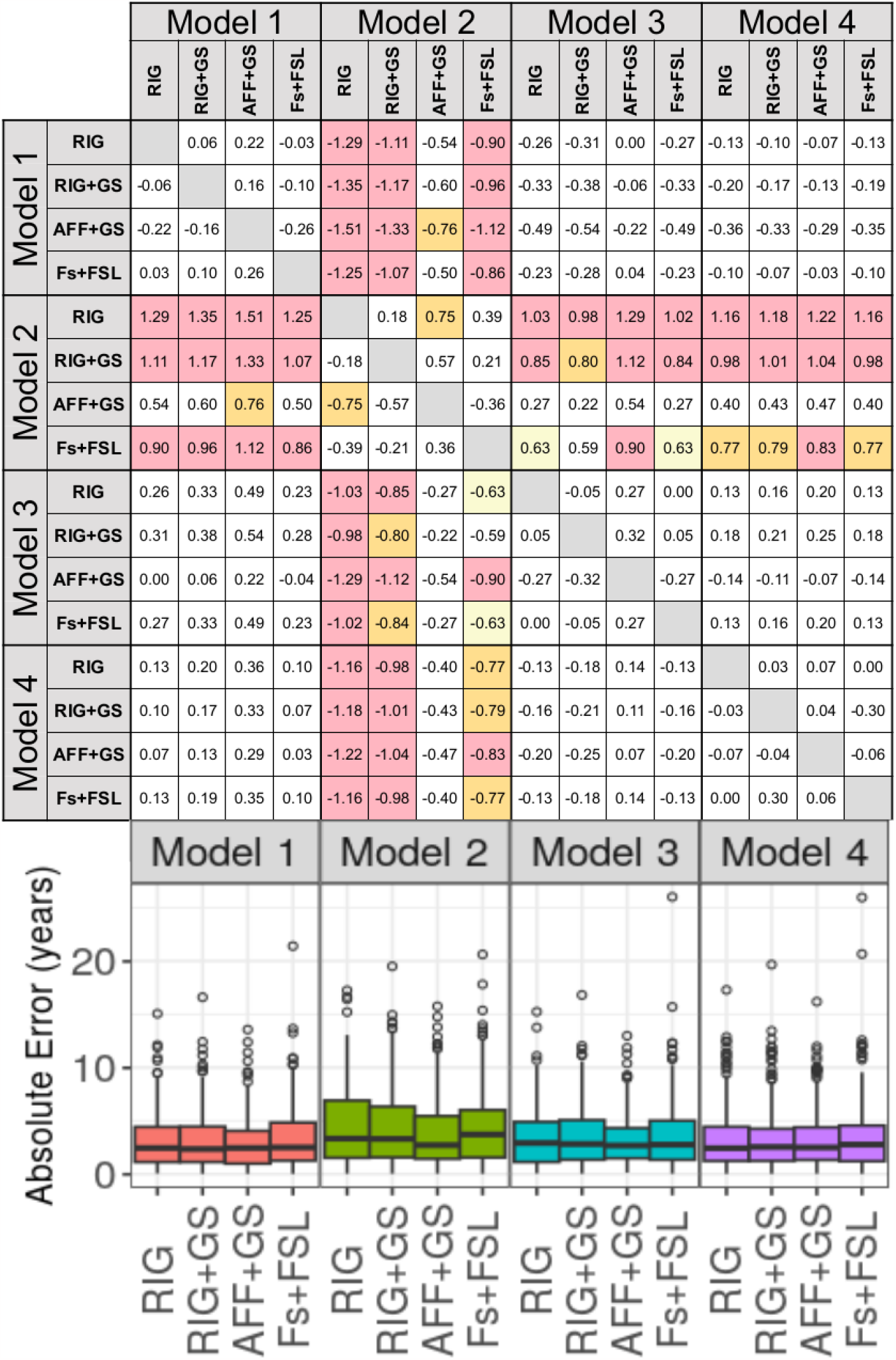
Results of LMEM post-hoc pairwise statistical tests for all MRI preprocessing and model architecture combinations. The color of each square marks statistical significance: red for *p* < 0.001, orange for 0.01 ≤ *p* < 0.001, yellow for 0.05 ≤ *p* < 0.01 and white for *p* > 0.05 (not significant).

In terms of sensitivity to preprocessing, Model 2 again displayed the most variability. Conversely, the 3D models demonstrated a more stable performance, with none experiencing an increase in MAE exceeding 0.32 years. Particularly, Model 4 showed notable robustness to the change in preprocessing, registering a MAE fluctuation within a range of 0.07 years (cf. Table 1).

Compared to the RIG pipeline, the RIG+GS included gray scale correction steps, however, resulted in only a marginal overall decrease in MAE. None of the differences were statistically significant (cf. Figure 4). When switching between rigid (RIG+GS) and affine registration (AFF+GS), we observed an improvement in performance for all models; for instance, as much as 0.57 years for Model 2. With Model 1, the reduction of MAE was 0.16 years to 2.96 years, which was the best MAE score reported in this study. While each model showcased enhanced performance with datasets AFF+GS, this improvement was statistically significant only for Model 2. Specifically, the difference was significant between AFF+GS and RIG preprocessing pipeline (*p* = 0.003). Interestingly, all models trained on the Fs+SL preprocessing presented higher MAE scores than those trained using theAFF+GS pipeline. This occurred despite both methods including registration and grayscale corrections. Even though the differences weren’t statistically significant, this observation suggests that the choice of software might still play a pivotal role in the outcomes.

### 4.3. Performance on unseen data

In this experiment, we evaluated the performance of 16 model ensembles on unseen data. In general, new data may come from a different MRI scanner or have undergone different preprocessing than the data used to train the models. In this experiment, we assumed that the unseen data had been preprocessed in the same way as the training data, which is a common scenario in practice.

To evaluate the performance of the models on the unseen data, we predicted brain age for all 1493 T1w scans of the UKB dataset without any additional training. For each model and preprocessing combination, we again averaged the results across five pretrained models with different weight initializations. This resulted in a total of 16 predictions for each T1w image, which served as our baseline.

By inspecting the Supplementary Figure 9, we observed a systematic offset of age prediction across the whole age span. This offset was inherent to each combination of architecture and preprocessing prediction and can be reduced by applying an offset correction (cf. Section 3.4). We comparatively evaluated *1)* the baseline predictions of uncorrected mean ensemble, and *2)* the offset corrected predictions, computed by deducting the ME from the predicted brain age value.

For estimating the influence of preprocessing pipeline on model performance on unseen data we fit a LMEM model with architecture, preprocessing, presence or absence of offset correction, their two-way and three-way interactions as fixed effects, and subject ID as random effect. ANOVA test confirmed that this model explained more variability than the model with no interactions (*p* < 0.001) and the model with two-way interactions (*p* < 0.001). ANOVA test of effects show all main effects, their two-way, and three-way interactions as statistically significant (*p* < 0.001). Detailed results of LMEM model and ANOVA test are presented in the Supplementary Table 8.

Table 2 show the mean ensemble MAE values for the 16 combinations of preprocessing and model architecture. The baseline MAE values range from 5.26 years for the 2D Model 2 with RIG+GS preprocessing to 3.33 years for Model 1 with Fs+FSL preprocessing. Bias, as measured by the ME shows, that all models on average underestimate brain age, when used on new dataset. Notably, the 2D model consistently exhibited the smallest bias, however, it also displayed the largest standard deviation in error across all preprocessing pipelines by roughly one year.

**Table 2:** MAE on unseen dataset (UKB), preprocessed in the same manner as multi-site training data. Results are presented for 16 models and preprocessing combinations, with and without additional offset correction. Best MAE values wrt. model architecture (*rows*) are <u>underlined</u>, while best values wrt. image preprocessing procedure (*columns*) are marked in **bold**. All numbers are in years.

|  | Corr. | RIG |  | RIG+GS |  | AFF+GS |  | Fs+FSL |  |
| --- | --- | --- | --- | --- | --- | --- | --- | --- | --- |
|  |  | ME (sd) | MAE (sd) | ME (sd) | MAE (sd) | ME (sd) | MAE (sd) | ME (sd) | MAE (sd) |
| Model 1 | none | -3.65 ± 4.13 | 4.48 ± 3.22 | -3.60 ± 4.19 | 4.47 ± 3.25 | -2.10 ± 4.21 | 3.73 ± 2.88 | -0.73 ± 4.09 | **3.33 ± 2.48** |
|  | offset | 0.0 ± 4.13 | **3.26 ± 2.54** | -0.0 ± 4.19 | 3.33 ± 2.55 | -0.0 ± 4.21 | 3.31 ± 2.60 | -0.0 ± 4.09 | 3.28 ± 2.43 |
| Model 2 | none | -1.70 ± 6.26 | 5.07 ± 4.05 | -1.65 ± 6.08 | 4.90 ± 3.94 | -1.58 ± 5.53 | **4.32 ± 3.78** | 0.03 ± 5.74 | 4.54 ± 3.52 |
|  | offset | 0.0 ± 6.26 | 4.95 ± 3.84 | -0.0 ± 6.08 | 4.75 ± 3.79 | 0.0 ± 5.53 | **4.21 ± 3.58** | -0.0 ± 5.74 | 4.54 ± 3.52 |
| Model 3 | none | -3.64 ± 4.56 | 4.73 ± 3.42 | -4.38 ± 4.64 | 5.26 ± 3.61 | -2.26 ± 4.51 | 3.93 ± 3.16 | -2.00 ± 4.32 | **3.78 ± 2.90** |
|  | offset | -0.0 ± 4.56 | 3.61 ± 2.79 | 0.0 ± 4.64 | 3.70 ± 2.79 | -0.0 ± 4.51 | 3.51 ± 2.82 | -0.0 ± 4.32 | **3.42 ± 2.64** |
| Model 4 | none | -4.42 ± 4.69 | 5.19 ± 3.83 | -3.29 ± 4.69 | 4.49 ± 3.55 | -1.64 ± 4.4 | 3.65 ± 2.95 | -0.71 ± 4.57 | **3.63 ± 2.86** |
|  | offset | 0.0 ± 4.69 | 3.69 ± 2.90 | -0.0 ± 4.69 | 3.71 ± 2.86 | 0.0 ± 4.40 | **3.45 ± 2.73** | 0.0 ± 4.57 | 3.59 ± 2.83 |

Figure 5 shows the pairwise difference in marginal means and their statistical significance between the preprocessing procedures, conditional on the model architecture and the presence or absence of offset correction, for the above-mentioned LMEM. Prior to offset correction, datasets with affine correction consistently demonstrated the best performance. Specifically, for Model 1, the results from the Fs+FSL dataset outperformed those from all other preprocessing pipelines. For Models 2, 3, and 4, both Fs+FSL and AFF+GS datasets showed superior performance compared to the rest of the preprocessing pipelines. However, there was no significant distinction between the two.

**Figure 5.**
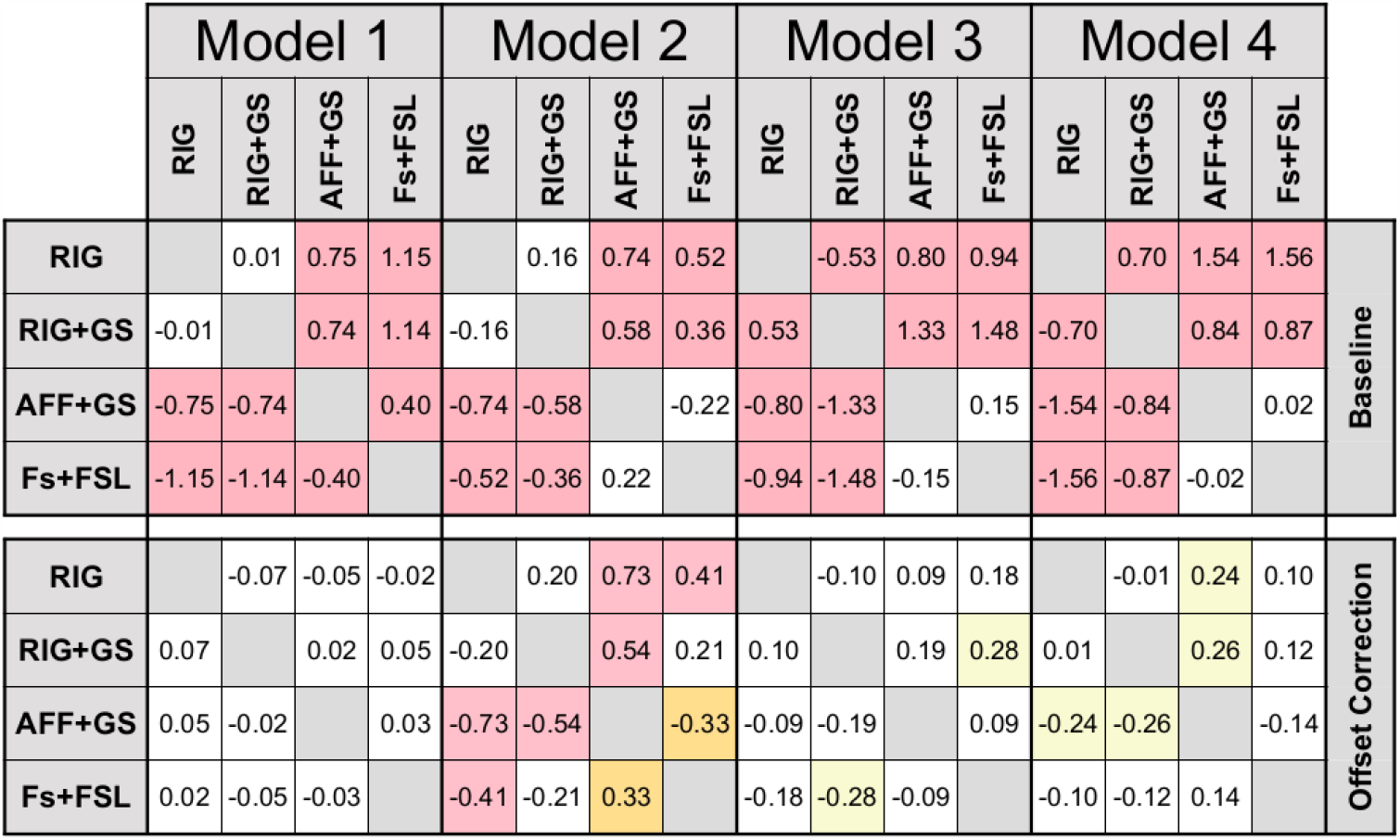
The pairwise difference in marginal means between preprocessing procedures conditional on model architecture and the presence or absence of offset correction. The preprocessing procedure of test data was the same as the preprocessing procedure applied to the train data. The color of each square marks the significance level of difference: red for *p* < 0.001, orange for 0.01 ≤ *p* < 0.001, yellow for 0.05 ≤ *p* < 0.01 and white for *p* > 0.05 (not significant).

When comparing model architectures, Models 1 and 4 consistently outperformed Models 2 and 3, trained on reduced information. The only exception was with the RIG preprocessing procedure, where Model 1 alone excelled. Even following offset correction, the 3D models maintained a performance edge over the 2D model.

Applying offset correction reduced MAE by 0.54 years on average. The distinction in performance between RIG and RIG+GS remained non-significant for all models (cf. Figure 5).Model 1 yielded the best performance across all preprocessing pipelines, achieving an overall best MAE of 3.31 ± 2.60 with the RIG preprocessing. Although the superior results from RIG might be surprising, it’s critical to note that Model 1 demonstrated robustness to change in preprocessing, exhibiting MAE within the range of 0.07 years. Model 2 was the most sensitive to preprocessing. It performed best when trained with affine registration, indicating its sensitivity to spatial information.

### 4.4. Performance on unseen data with new image preprocessing

We further considered the cumulative effect of dataset not used during model training, additionally with different image preprocessing as dataset used during model training. The UKB was preprocessed by the dataset provider, described in Section 3.2.1. Without additional training we predicted the age for all 80 trained models. The model predictions were ensembled across five models with different weight initialization, which resulted in 16 predictions per each T1w MRI (baseline). As in the previous experiment, we comparatively evaluated *1)* the baseline predictions and *2)* the offset-corrected predictions (cf. Section 3.4). The prediction offset on the new dataset is constant across the whole age span, as evidenced by Supplementary Figure 9.

The MAE and ME metrics of the 16 mean ensembles are presented in Table 3. For estimating the influence of preprocessing pipeline, we fit a LMEM model with architecture, preprocessing, presence or absence of offset correction, their two-way and three-way interactions as fixed effects, and subject ID as random effect. ANOVA test confirmed that this model explained more variability than simpler LMEM models (*p* < 0.001). ANOVA test of effects show all main effects, their two way, and three way interactions as statistically significant (*p* < 0.001). Details are provided in Supplementary Table 9.

**Table 3:**
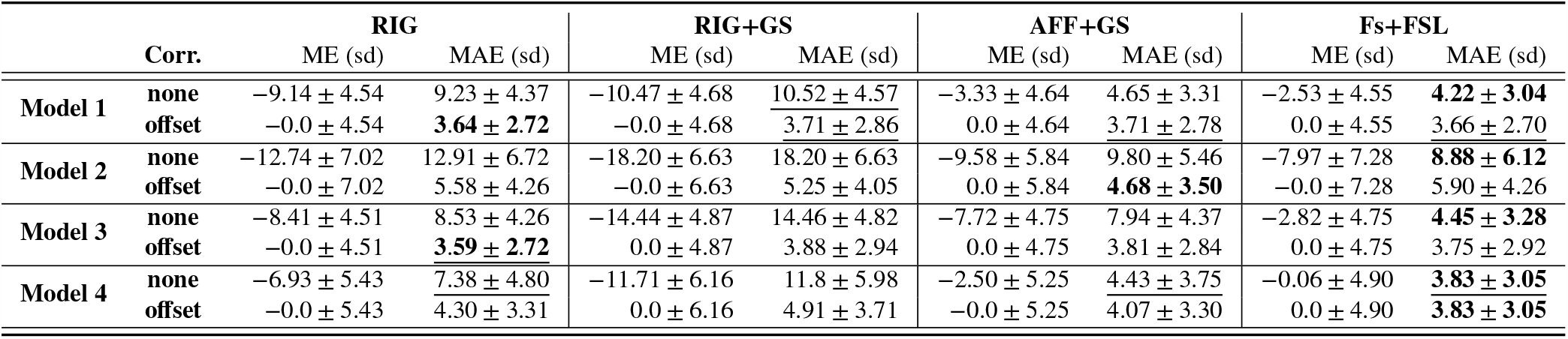
Mean ensemble MAE values for 16 preprocessing and model architecture combinations, with and without offset correction on UKB dataset with new preprocessing procedure. Best MAE values wrt. model architecture (*rows*) are <u>underlined</u>, while best values wrt. image preprocessing procedure (*columns*) are marked in **bold**. All numbers are in years.

|  | Corr. | RIG |  | RIG+GS |  | AFF+GS |  | Fs+FsL |  |
| --- | --- | --- | --- | --- | --- | --- | --- | --- | --- |
|  |  | ME (sd) | MAE (sd) | ME (sd) | MAE (sd) | ME (sd) | MAE (sd) | ME (sd) | MAE (sd) |
| Model 1 | none | -9.14 ± 4.54 | 9.23 ± 4.37 | -10.47 ± 4.68 | <u>10.52 ± 4.57</u> | -3.33 ± 4.64 | 4.65 ± 3.31 | -2.53 ± 4.55 | **4.22 ± 3.04** |
|  | offset | -0.0 ± 4.54 | **3.64 ± 2.72** | -0.0 ± 4.68 | <u>3.71 ± 2.86</u> | 0.0 ± 4.64 | 3.71 ± 2.78 | 0.0 ± 4.55 | <u>3.66 ± 2.70</u> |
| Model 2 | none | -12.74 ± 7.02 | 12.91 ± 6.72 | -18.20 ± 6.63 | 18.20 ± 6.63 | -9.58 ± 5.84 | 9.80 ± 5.46 | -7.97 ± 7.28 | **8.88 ± 6.12** |
|  | offset | -0.0 ± 7.02 | 5.58 ± 4.26 | -0.0 ± 6.63 | 5.25 ± 4.05 | 0.0 ± 5.84 | **4.68 ± 3.50** | -0.0 ± 7.28 | 5.90 ± 4.26 |
| Model 3 | none | -8.41 ± 4.51 | 8.53 ± 4.26 | -14.44 ± 4.87 | 14.46 ± 4.82 | -7.72 ± 4.75 | 7.94 ± 4.37 | -2.82 ± 4.75 | **4.45 ± 3.28** |
|  | offset | -0.0 ± 4.51 | **3.59 ± 2.72** | 0.0 ± 4.87 | 3.88 ± 2.94 | 0.0 ± 4.75 | 3.81 ± 2.84 | 0.0 ± 4.75 | 3.75 ± 2.92 |
| Model 4 | none | -6.93 ± 5.43 | <u>7.38 ± 4.80</u> | -11.71 ± 6.16 | 11.8 ± 5.98 | -2.50 ± 5.25 | <u>4.43 ± 3.75</u> | -0.06 ± 4.90 | **3.83 ± 3.05** |
|  | offset | -0.0 ± 5.43 | 4.30 ± 3.31 | 0.0 ± 6.16 | 4.91 ± 3.71 | -0.0 ± 5.25 | 4.07 ± 3.30 | 0.0 ± 4.90 | **3.83 ± 3.05** |

**Table 4:** Age statistics, i.e. span, mean age (***μ***_**age**_) and associated standard deviation (**sd**_**age**_) in *years*, per dataset of included T1w subject scans in train, test and validation datasets (*top*) and in the new unseen site and test-retest datasets (*bottom*).

| Dataset | N<sub>scans</sub> | Age span | $\mu_{\text{age}} \pm \text{sd}_{\text{age}}$ |
| --- | --- | --- | --- |
| ABIDE I<sup>8</sup> | 161 | 18.0 – 48.0 | 25.7 ± 6.4 |
| ADNI<sup>9</sup> | 248 | 60.0 – 90.0 | 76.2 ± 5.1 |
| CamCAN (Shafto et al., 2014; Taylor et al., 2017)<sup>10</sup> | 624 | 18.0 – 88.0 | 54.2 ± 18.4 |
| CC-359 (Souza et al., 2018)<sup>11</sup> | 349 | 29.0 – 80.0 | 53.5 ± 7.8 |
| FCON 1000<sup>12</sup> | 572 | 18.0 – 85.0 | 45.3 ± 18.9 |
| IXI<sup>13</sup> | 472 | 20.1 – 86.2 | 49.0 ± 16.2 |
| OASIS-2 (Marcus et al., 2010)<sup>14</sup> | 78 | 60.0 – 95.0 | 75.6 ± 8.4 |
| **Total** | 2504 | 18.0 – 95.0 | 52.1 ± 19.1 |

*Aim: Test (Unseen T1w scans)*
| Dataset | N<sub>subj</sub> | Age span | $\mu_{\text{age}} \pm \text{sd}_{\text{age}}$ |
| --- | --- | --- | --- |
| UK Biobank (Miller et al., 2016) | 1493 | 48.5 – 80.4 | 63.1 ± 7.2 |

<sup>1</sup> Data available at: http://fcon_1000.projects.nitrc.org/indi/abide/abide_I.html
<sup>2</sup> Data available at: http://adni.loni.usc.edu/
<sup>3</sup> Data available at: https://camcan-archive.mrc-cbu.cam.ac.uk/dataaccess/
<sup>4</sup> Data available at: https://sites.google.com/view/calgary-campinas-dataset/download
<sup>5</sup> Data available at: http://fcon_1000.projects.nitrc.org/indi/enhanced/neurodata.html
<sup>6</sup> Data available at: https://brain-development.org/ixi-dataset/
<sup>7</sup> Data available at: https://www.oasis-brains.org/

For baseline results, the predicted age is generally underestimated for the unseen UKB detaset, with ME as low as −18.20 years for Model 2 on RIG+GS dataset, and only −0.06 years for Model 4 with Fs+FSL preprocessing. This may be expected considering the smaller age span of the UKB population versus the multi-site train set population (cf. Supplementary Table 4). Large AE values close to 40 years were observed for Model 2. The model exhibits a large bias and errors across all four image preprocessing procedures with MAE ranging from 18.20 years for RIG+GS preprocessing to 8.88 years for Fs+FSL preprocessing. Additionally, the variance of predictions is larger than for 3D models, as seen from Supplementary Figure 10.

Figure 6 displays the pairwise differences in marginal means of preprocessing, conditional on the model architecture, and the presence or absence of offset correction. For the baseline results, large and statistically significant differences in performance were shown between all combinations of the preprocessing pipelines and models. The disparity is performance is most pronounced between models trained on datasets with affine registration and those with rigid registration; for the former, the MAE nearly doubled. For instance, Model 1, when trained on the RIG+GS dataset, yielded an MAE of 10.52 years, whereas it was 4.22 years with the Fs+FSL preprocessing. For all models, the best baseline result was achieved on Fs+FSL, which can be attributed to the fact that the same software was used for preprocessing of UKB dataset (Section 3.2.1).

**Figure 6.**
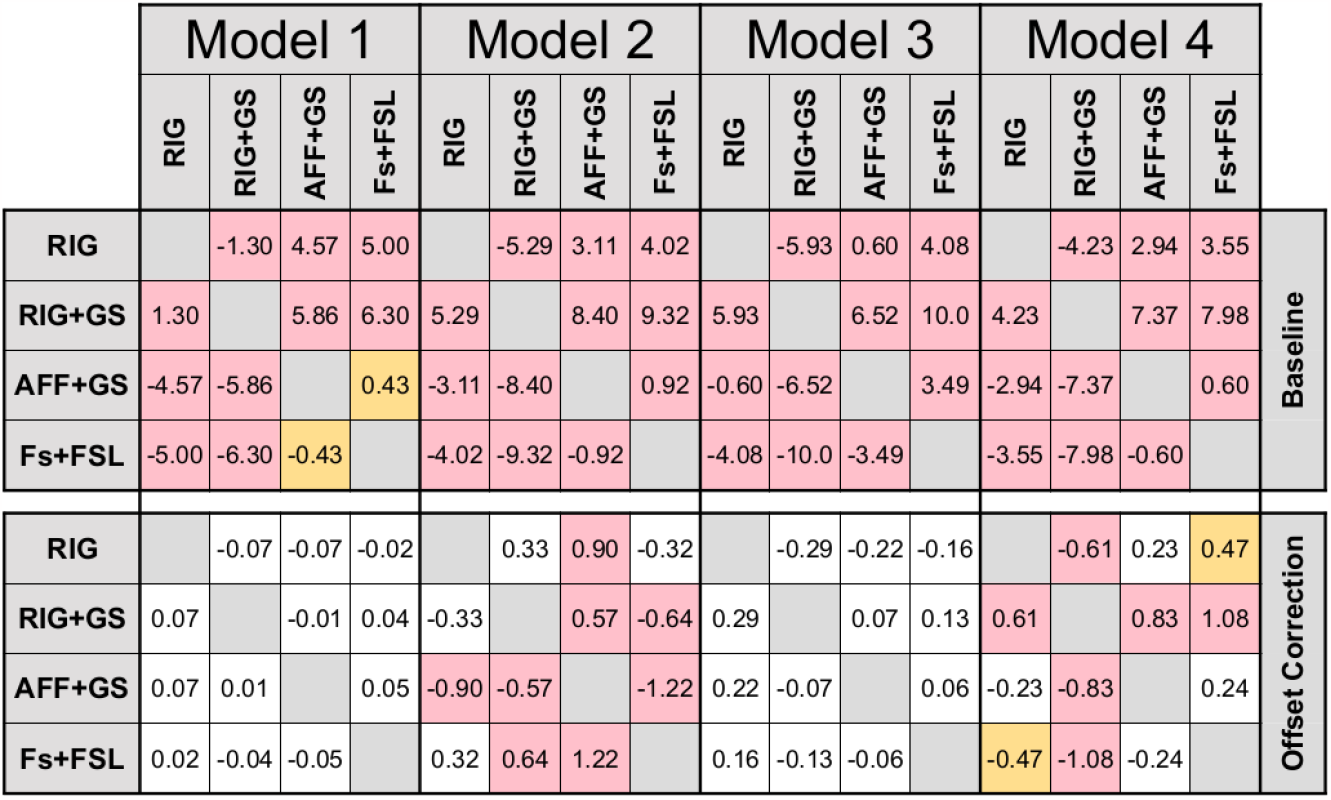
The pairwise difference in marginal means between preprocessing procedures conditional on model architecture. The preprocessing procedure of test data differed from preprocessing procedure of train data. The color of each square marks the significance of difference: red for *p* < 0.001, orange for 0.01 ≤ *p* < 0.001, yellow for 0.05 ≤ *p* < 0.01 and white for *p* > 0.05 (not significant).

**Figure 7.**
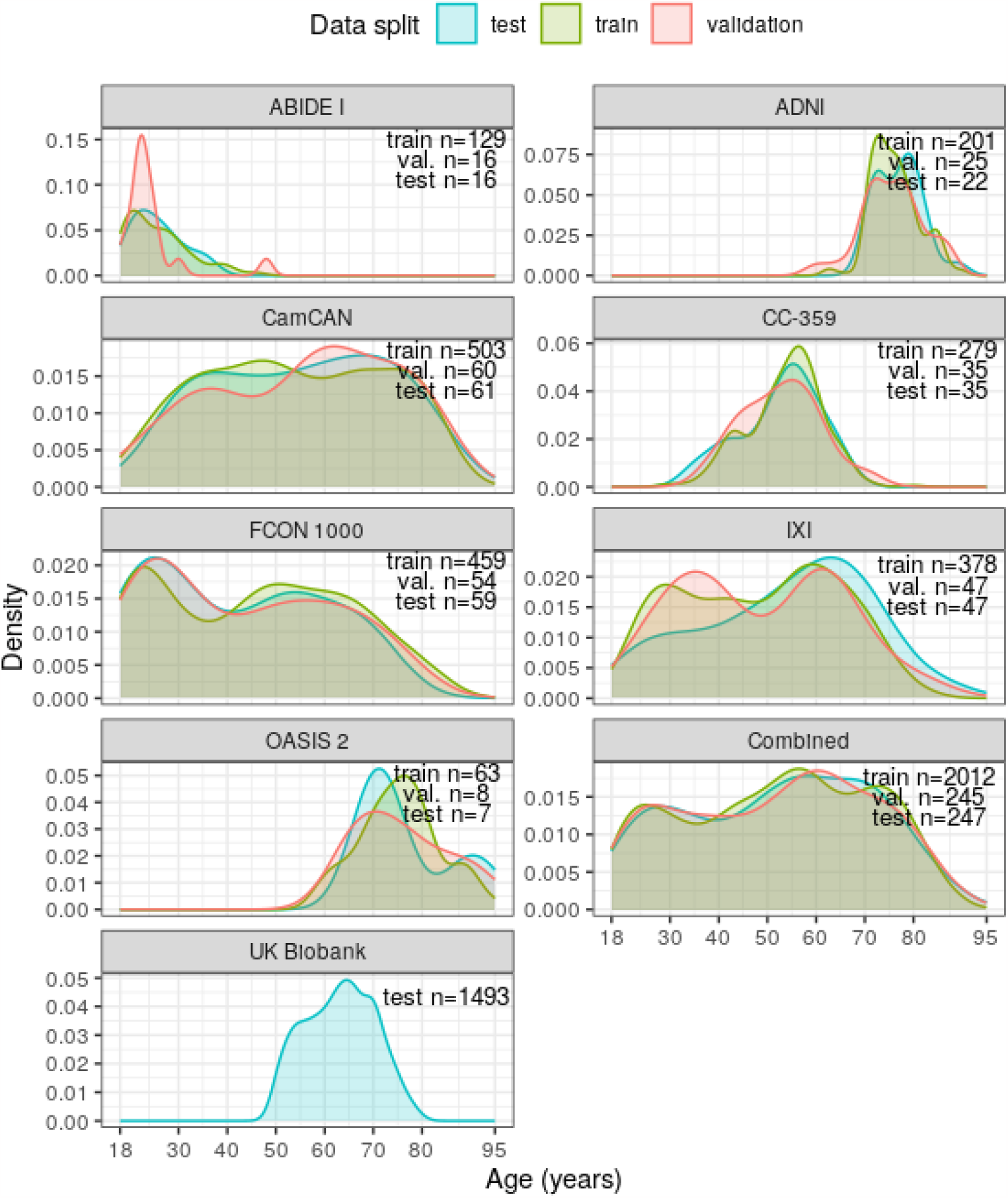
Density of age distribution per each dataset and combined multi-site dataset, depicted for train, test and validation set splits.

**Figure 8.**
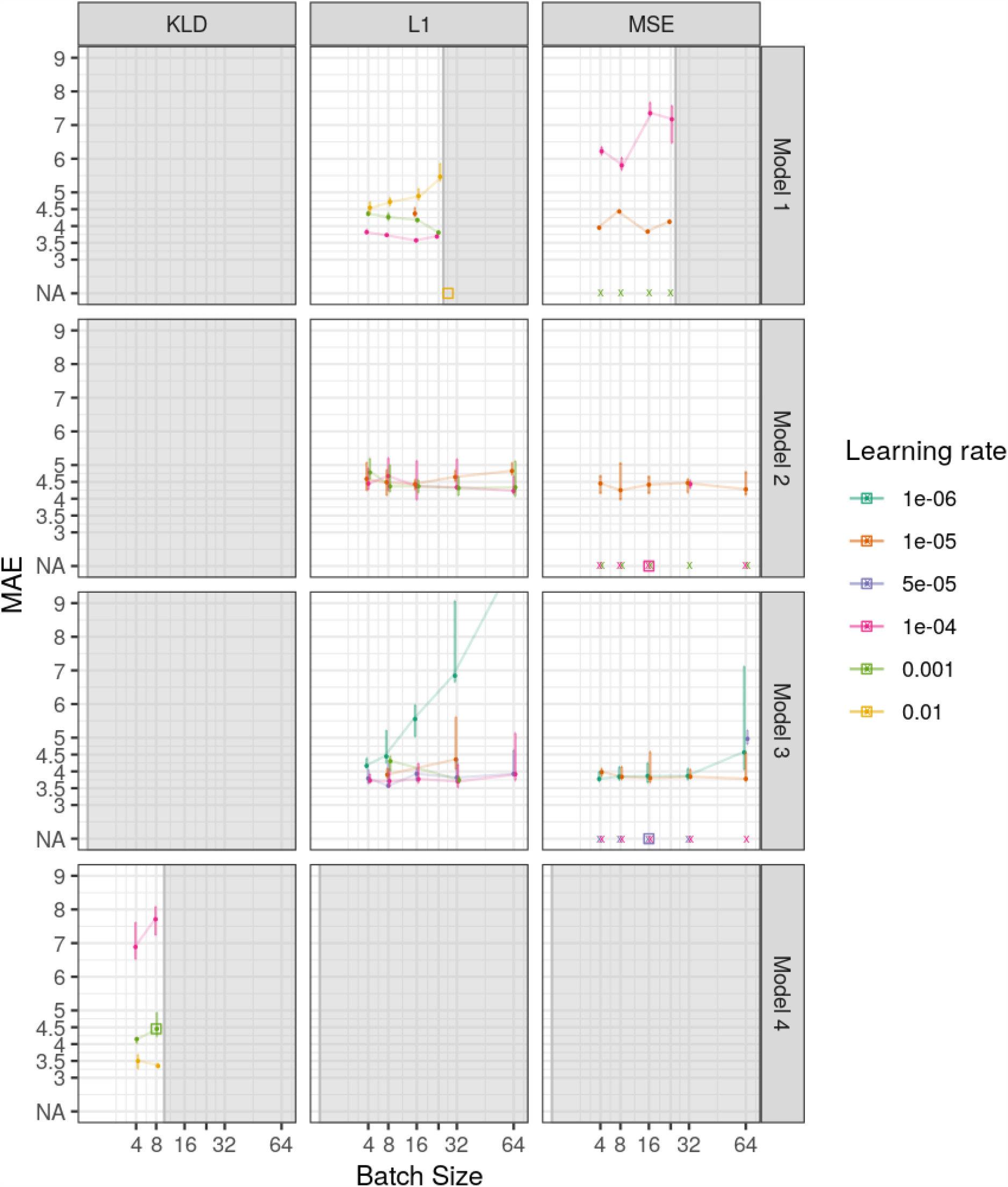
Median, minimal and maximal MAE value of 10 last training epochs for each hyperparameter setting. The hyperparameter values proposed in original research of four models are marked with *square*, the ones resulting in training divergence are *marked as NA and with a cross*. Hyperparameter space for large batch sizes was inaccessible due to hardware limitations and is *grayed out*.

**Figure 9.**
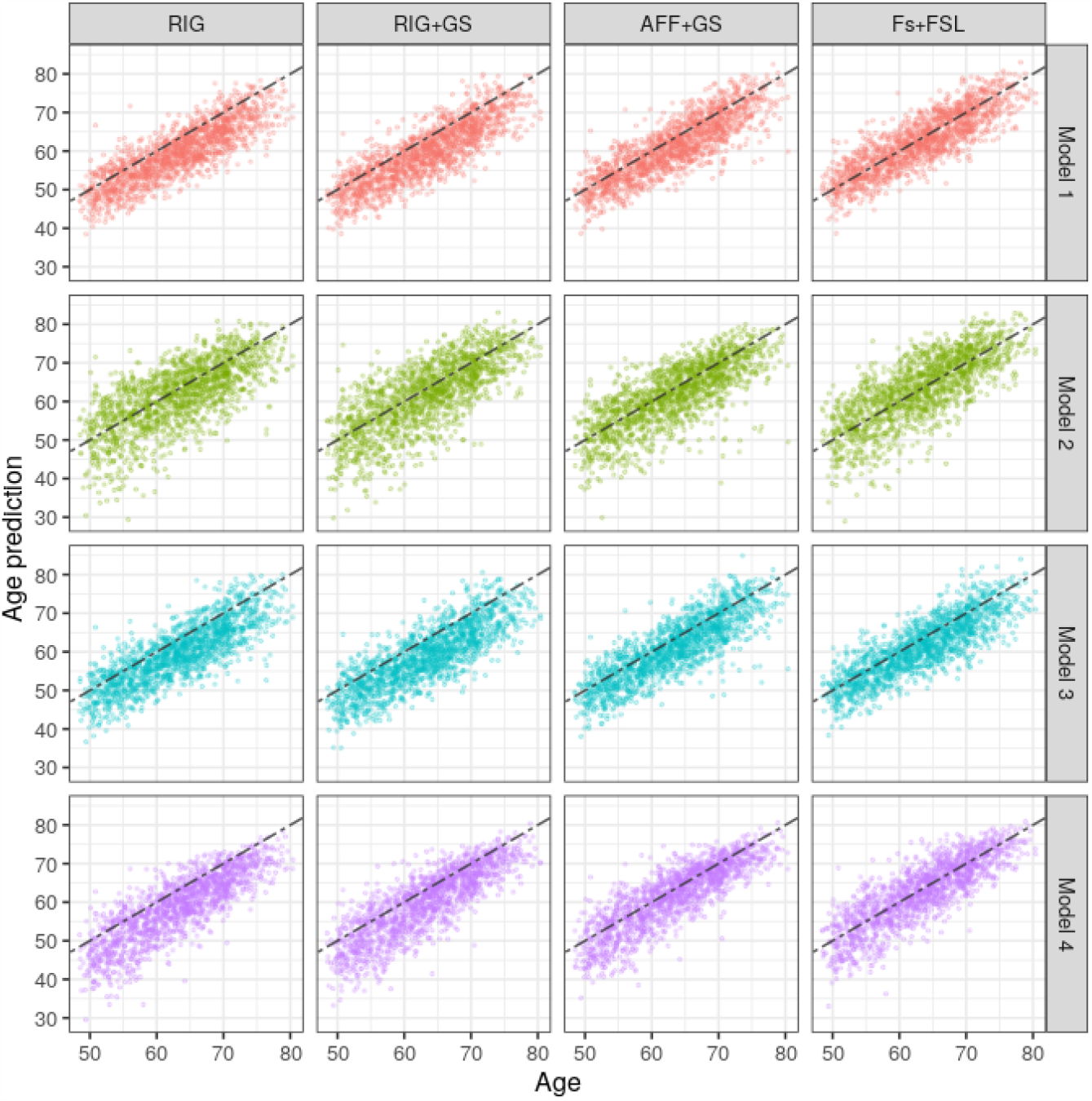
Model predictions on UKB dataset, preprocessed in the same manner as the training set.

**Figure 10.**
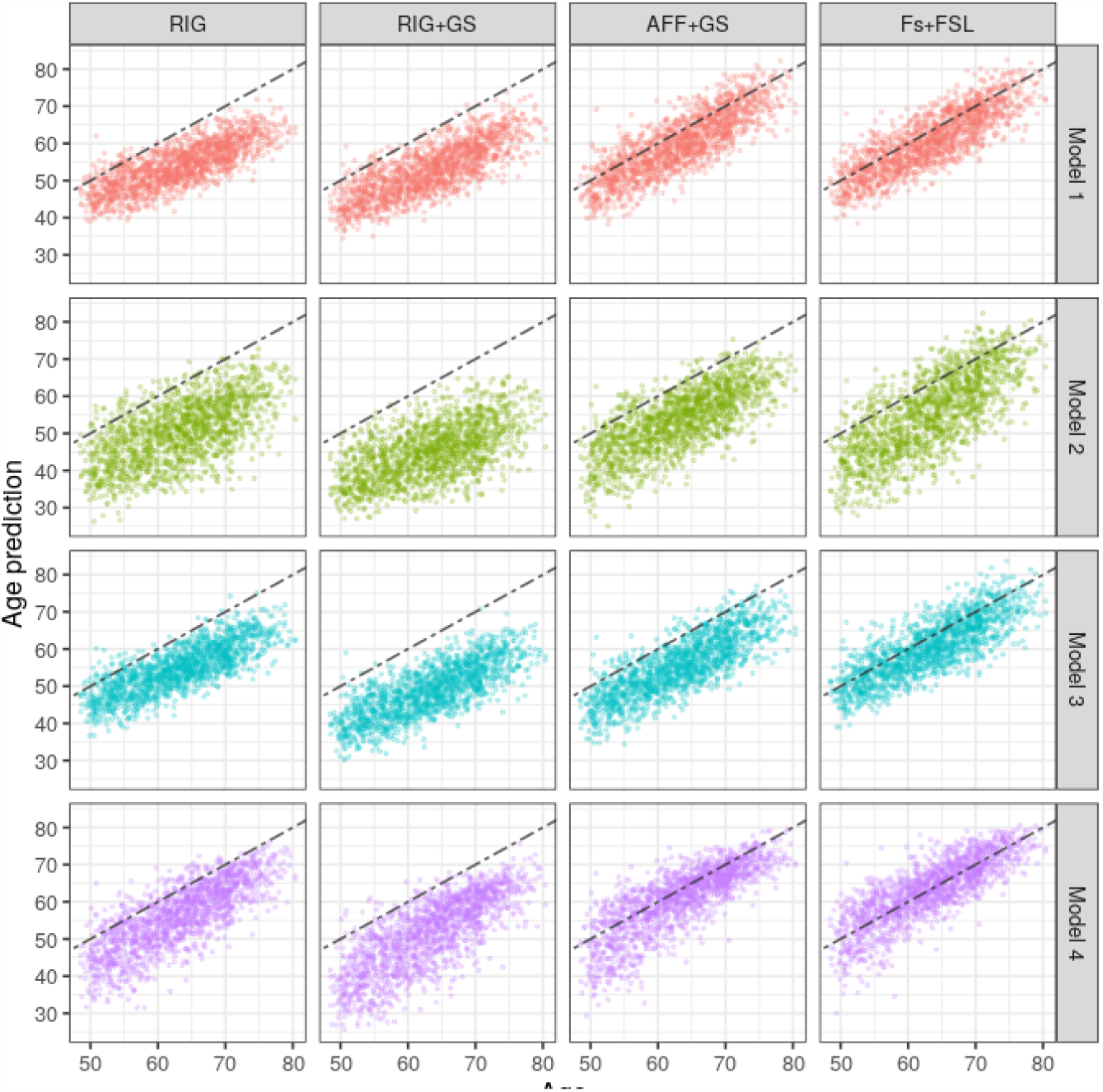
Model predictions on UKB dataset, preprocessed using different preprocessing pipeline as the training set.

Correction of systematic offset improved MAE for 4.56 years on average. Despite offset correction, the introduction of different preprocessing of the test set, led to the average increase in MAE 0.5 years, when compared to the findings to Table 2 in Section 4.3. This increase was smallest for model Model 3 and largest for Model 4. Additionally, the standard deviation of absolute error increased by up to 0.85 years, and the standard deviation of error also grew by as much as 1.5 years.

On offset corrected results, Model 1 generally exhibited best performance, only being surpassed by Model 3 on the RIG preprocessing, achieving an overall best MAE of 3.59 years. For these two architectures, there were no significant differences in performance based on the preprocessing procedure used on training data (cf. Figure 6). Despite the change in datasets and their preprocessing between training and testing phases, Model 1 again showcased its robustness, displaying an MAE variation of only 0.07 years. While Model 4 displayed optimal performance prior to offset correction, it only matched Model 1 on the Fs+FSL dataset after applying the offset correction, hinting at its susceptibility to the change in preprocessing procedures between training and test datasets. As before, Model 2 was outperformed by all three 3D models. Nevertheless, the introduction of affine registration enhanced Model 2’s performance.

## 5. Discussion

This work studied the effect of four different T1w preprocessing procedures and implementations on the brain age prediction accuracy using deep learning-based models. For this purpose we implemented, trained and evaluated four CNN architectures presented in the brain age literature. Each model was initialized and trained five times and we reported the mean values of the five model predictions across all model architecture and T1w preprocessing combinations.

For model training, we compiled a multi-site dataset sourced from seven public repositories. These datasets encompassed images from a range of MRI vendors, spanning both 1.5 T and 3 T field strengths. We further considered predicting on a new T1w image dataset, not seen during model training, which was preprocessed the same as the training dataset, or in a different manner using different operators or software implementation.

The use of the multi-site dataset for model training generally increases the model’s accuracy, since using a single-site training and test dataset, and/or dataset with limited age span, may lead to overoptimistic accuracy of predictions, which cannot be reproduced when applying such a model to an unseen (multi-site) dataset. We have experimentally verified this claim (results not shown) by assembling a training dataset from the UKB subjects and then trained and evaluated our Model 1 on the (age span adjusted) multi-site and UKB test datasets. On the latter we achieved a MAE of 2.28 ± 1.85 years, which, however, could not be reproduced on the multi-site test dataset (MAE was 3.39 ± 2.38), even after offset correction. Contrarily, when using the multi-site dataset for training, the obtained MAE can be surprisingly well reproduced on the unseen (UKB) data (MAE was 3.33 ± 2.49). The aforementioned experiment clearly underscored the need to use multi-site train and test datasets in order to ensure reproducibility and validity of the findings.

The point estimates of brain age accuracy like the MAE, which are usually reported in brain age literature, need to be statistically evaluated to enable one to draw generalizing conclusions. For this purpose we used the linear mixed-effects models (LMEMs), as they allow us to account for repeated measures on a subject level by including the subject ID as a random effect. As our results show, despite observed difference in point estimates of the MAE values, the difference may not be statistically significant. For instance, when comparing the performances of Models 1 and 3 obtained with the AFF+GS preprocessing (cf. Table 1), the seemingly relevant difference in MAE values of about 0.2 years was not statistically significant (cf. Figure 4).

### 5.1. Impact of T1w image preprocessing and model architecture

When comparing the effect of T1w preprocessing with respect to the four T1w preprocessing pipelines, a slightly higher brain age prediction accuracy (i.e. low MAE) across all models was observed for the AFF+GS preprocessing pipeline. Interestingly, the inclusion of gray scale correction into the pipeline, such as denoising and intensity inhomogeneity correction did not improve MAE, but was needed for accurate (linear) image registration used in our preprocessing pipelines.

In contrast to the “raw” preprocessing pipeline applied by Cole et al. (2017), we incorporated denoising to mitigate non-informative high-frequency noise in the MRI acquisitions. The effect of denoising is minimal and primarily intended to suppress the non-informative high-frequency noise. Using ensembles of all four model architectures trained on RIG preprocessing, we predicted age on a multi-site test dataset and applied a LMEM with model architecture, denoising, and their interactions as primary factors (results not shown). Similar to our observations for other grayscale corrections, the ANOVA results of the main effect of denoising was not significant (*F* (1, 5712.8) = 0.025, *p* = 0.874), neither was the interaction effect between model architecture and denoising (*F* (3, 5712.0) = 0.410, *p* = 0.746).

Despite the considerable similarities between the T1w preprocessing pipelines, we observed a difference in performance between models trained using common software such as FSL and FreeSurfer and AFF+GS pipeline. This shows that even when training brain age prediction models on the same source data, but with different implementations of T1w preprocessing software, the obtained results may not be directly comparable.

Even comparisons with results from original papers, where the CNN models evaluated in this study were proposed, are not straightforward due to disparities in the size and age structure of the training datasets. Interestingly, the MAE of Model 1 reported herein was 1 year lower than the MAE reported by Cole et al. (2017), despite the fact that the T1w preprocessing (RIG), structure and size of training set were similar. We attribute the improvement partially to the mean ensembling and largely to extensive hyperparameter tuning.

In general, all four model architectures performed considerably better if T1w preprocessing involved linear (affine) registration, i.e. the AFF+GS and Fs+FSL. This indicates the importance of good spatial normalization of the input T1w scans, which eliminates the inter-subject variance due to head size differences and MRI-acquisition related geometric artifacts. However, pairwise comparison was only marginally statistically significant between the AFF+GS and RIG, and AFF+GS and Fs+FSL T1w preprocessing pipelines. This enhancement particularly benefited models receiving limited input information. Specifically, Model 3, which was trained on downsampled 3D images, and to a greater extent, Model 2, trained on 2D slices, which demonstrated sensitivity to spatial information across all experiments. Our results are in inline with the study by Peng et al. (2021), wherein the T1w preprocessing procedures including either linear or non-linear registration were compared, resulting in slight favor of the latter.

In their review Tanveer et al. (2023) discuss computational complexity and call on research community to further focus on the 2D CNN brain age prediction models. Our results show a statistically significant inferiority of the implemented 2D model versus all tested 3D models. The 2D model only performed on par, without statistically significant underperformance, when trained on the AFF+GS preprocessed dataset. This finding is in line with Feng et al. (2020), who showed that a 2D model, which is designed analogous to a 3D model, performs significantly worse. Therefore any future 2D implementations cannot be naive reimplementations of the 3D models, but need to introduce a methodological improvement, like for instance Jönemo et al. (2022), predicting age from 2D projections of the 3D MRI volumes.

The final aspect when comparing different preprocessing procedures is the computational complexity. To use the brain age as a prognostic biomarker in clinical practice, the T1w brain MRI should be processed in a reasonably short time. Despite the more extended execution times of some of the presented preprocessing pipelines (ranging from 1.5 to 16.5 minutes), substantial gains seem achievable with the utilization of GPU (re-)implementations. The lengthy training times of the deep learning models do not appear limiting, as they are carried out off-line. In contrast, the model inference time per brain age prediction is typically only a few seconds, making it negligible compared to the T1w preprocessing time.

However, in situations where resources are limited, there is a certain trade-off between model implementability and accuracy, as also pointed out by Dartora et al. (2023). When compute resources are limited and larger models cannot be trained, our results demonstrate a pivotal role of extensive preprocessing. While models trained on full resolution 3D data are less sensitive to preprocessing variations, extensive preprocessing improves accuracy especially for the smaller-footprint models trained on 2D or downsampled 3D MRI images. Our findings further underscore the importance of thorough preprocessing when predicting on new site datasets. In the absence of offset correction, extensive preprocessing acts as a form of data harmonisation, ensuring consistency of predictions across varying data sources. It is essential to consider both the computational demands/resources and the desired accuracy to identify the most suitable model-pipeline combination for a given research setting.

### 5.2. Performance on new unseen scanner data

In contrast to the differences in MAE predictions on multi site-dataset, where only marginally significant differences were observed, the differences between models and preprocessing procedures were significant when inferring on the new unseen data. Among the T1w preprocessing procedures evaluated, those that were more extensive (Fs+FSL and AFF+GS) exhibited the lowest brain age prediction errors when predicting on new site dataset (before offset correction) and were significantly better than those using only rigid registration for all models. The MAE marginal difference between Fs+FSL and AFF+GS was significant only for Model 1. After offset correction was applied, datasets preprocessed with affine registration continued to outperform, although Model 1 displayed equivalent performance across all preprocessing pipelines. Such findings show that extensive preprocessing also potentially serves as an instrument for data harmonization.

Despite recent efforts to use minimally preprocessed T1w images (Dartora et al., 2023; Fisch et al., 2021) and calls for further development of models on routine MRIs (Tanveer et al., 2023), our findings suggest that using more extensive T1w preprocessing can improve the prediction accuracy of brain age models even on datasets obtained from a new site. The sensitivity to the type of spatial registration of the T1w image to the brain atlas space was particularly crucial for the 2D model, which was trained on 15 axial slices and performed significantly better on datasets with the affine registration, resulting in an improvement in MAE of 0.5 years.

However, it is worth noting that other factors, such as the size and characteristics of the dataset, may also influence the brain age prediction accuracy. For example, before offset correction, the models tend to underestimate the age of the subjects in the UKB dataset, which may be due to the higher age of the individuals in the training dataset. Additionally, the observed MAE values of 2.96 years for the multi-site test dataset and 3.26 for the new-site dataset were comparable, which may be partially attributed to the smaller age range of the subjects in the UKB dataset. However, the MAE is unlikely to increase proportionally with the increase of age range for adult datasets as assumed by Cole et al. (2019). For instance, in an experiment conducted by Peng et al. (2021), Model 4 was trained on UKB and in a separate experiment on dataset with age ranging from 17 to 90 years of somewhat similar sizes (i.e. 2600 and 2200 subjects) and achieved MAE values of 2.76 years and 2.9 years, respectively.

### 5.3. Transferability of model on dataset with new preprocessing

Differences in the level of applied T1w preprocessing between the training and test set played a crucial role when predicting brain age on new unseen scanner data. For instance, the values of MAE obtained with RIG and/or RIG+GS pipelines were more than double the values of the MAE obtained with the Fs+FSL and/or AFF+GS pipelines. These results are in line with the observation by Cole et al. (2017), who found a substantially reduced between-scanner reliability for a model trained on minimally preprocessed T1w images.

The performance before offset correction of all models was best on Fs+FSL pipeline. We attribute this to the similarity of the T1w preprocessing pipeline (and software) applied to the UKB dataset, as well as generally observed better performance of the models trained with the Fs+FSL and/or AFF+GS pipelines. The MAE on Fs+FSL was only about 1 year worse than the MAE obtained when the same T1w preprocessing was applied prior to offset correction, however also the standard deviations of errors increased. This shows that while models can bridge some gap between the preprocessing of train and test set, both bias and variance increase.

Regardless of the T1w preprocessing differences on the new unseen dataset with different preprocessing, the increase in MAE for Models 1 and 4 was 4 and 3.5 year, respectively, even before applying offset correction. However, when focusing solely on the models trained on the AFF+GS and Fs+FSL pipelines, this increase was only 1.35 and 0.85 years. This was comparable to or lower than the magnitude of the observed increase in most related literature (Jonsson et al., 2019; Dufumier et al., 2021).

Despite the small difference in MAE values for some models, most would be unusable in practice without applying offset correction. Therefore, the dataset dependent systematic bias should be mitigated by applying offset correction on (a subset of) the new unseen data.

After applying offset correction, we observed a similar pattern as when the preprocessing pipeline between training and testing sets remained unchanged. Model 1 consistently showed comparable performance, with minimal variations in MAE. In contrast, Model 2 was the most sensitive to the change in preprocessing procedure of the test set. Notably, Model 3, trained on downsampled images, achieved the best results, albeit by a slight margin. While it wasn’t the most consistent performer across all datasets, this shows that a robust 3D model trained on downsampled images can yield results on par with models trained on full-resolution 3D MRIs.

### 5.4. Impact of T1w image quality

The quality of the image plays a pivotal role in influencing brain age predictions. Our analysis showed that subpar image quality can lead to a substantial increase in MAE for all models and preprocessing pipelines, with some preprocessing procedures experiencing more than a twofold increase (results not shown). Among the various procedures, the Fs+FSL pipeline showed the most resilience to images of inferior quality. However, the prediction accuracy was still rather poor. This relative advantage is likely misleading due to the intensity normalization step in the Fs+FSL, which standardizes the mean intensity of the white matter, in turn reducing the Coefficient of Joint Variation and Contrast to Noise ratio. Nevertheless, further research is essential to fully understand and mitigate these impacts.

### 5.5. Note on reproducibility

The standardized dataset included multi-site train, validation and test T1w scans of 2504 healthy subjects in the span of 18 to 95 years, and test sets with new site T1w MRIs (*N* = 1493). All T1w MRI scans used in the study were obtained from public data sources^7^ and were subject to a strict visual quality check to eliminate poor quality scans or scans with failed T1w preprocessing.

In order to enable full reproducibility of the results of this study the lists of included subject IDs and the exact dataset split assignments as used in this study are provided in the Supplementary materials, while the implementations and dependencies of the T1w preprocessing routines, brain age models, scripts to re-run the experiments and carry out the performance evaluations and statistical analyses are disclosed at the public GitHub repository https://github.com/AralRalud/BrainAgePreprocessing.

## 6. Conclusion

In this paper we studied the effect of preprocessing procedure of T1w MRIs on the prediction accuracy of deep brain age models. We considered four preprocessing pipelines, which differed in the degree of freedom of T1w to brain atlas registration, the level of gray scale corrections and software implementations used. Our results for four different CNN architecture show that the choice of software implementation resulted in statistically significant increase in MAE, up to 0.75 years for the same model and dataset. We further show that applying the grayscale corrections does not significantly improve MAE of model predictions. The type of registration was shown to statistically significantly improve MAE when using affine compared to the rigid registration. Models trained on images with isotropic 1 × 1 × 1 *mm*^3^ spacing were less sensitive to the type of T1w preprocessing than the 2D model or the model trained on downsampled 3D images. Most affected by the (mis)registration of the input T1w MRI was the 2D model, since it was limited to only 15 axial slices, predefined in the MNI brain atlas space. In this case, the affine registration was crucial for good model performance, especially when predicting brain age on new dataset not seen during model training. Despite assumptions that models trained on less processed data are better suited for brain age prediction on new scanner datasets, not seen in model training, our results show that extensive T1w preprocessing in fact improves the generalization of brain age models when applied on new unseen datasets. Regardless of the model or the T1w preprocessing used, offset correction should be applied whenever predicting brain age on a new dataset with either the same or different T1w preprocessing as the one used in model training.

## Acknowledgments

Data collection and sharing for this project was partially provided by:

- **Alzheimer’s Disease Neuroimaging Initiative (ADNI) database (adni.loni.usc.edu)**. The investigators within the ADNI contributed to the design and implementation of ADNI and/or provided data but did not participate in analysis or writing of this report. A complete listing of ADNI investigators can be found at http://adni.loni.usc.edu/wp-content/uploads/how_to_apply/ADNI_Acknowledgement_List.pdf.
- **Cambridge Centre for Ageing and Neuroscience (CamCAN)**. CamCAN funding was provided by the UK Biotechnology and Biological Sciences Research Council (grant number BB/H008217/1), together with support from the UK Medical Research Council and University of Cambridge, UK.
- **OASIS Longitudinal**. Principal Investigators: D. Marcus, R, Buckner, J. Csernansky, J. Morris; P50 AG05681, P01 AG03991, P01 AG026276, R01 AG021910, P20 MH071616, U24 RR021382.
- **ABIDE I**. Primary support for the work by Adriana Di Martino was provided by the (NIMH K23MH087770) and the Leon Levy Foundation. Primary support for the work by Michael P. Milham and the INDI team was provided by gifts from Joseph P. Healy and the Stavros Niarchos Foundation to the Child Mind Institute, as well as by an NIMH award to MPM (NIMH R03MH096321).
- **the UK Biobank Resource** under Application Number 68981.

## A. Dataset and model details

### A.1. Dataset details

### A.2. Hyperparameter tuning and selection of loss function

Supplementary Figure 8 presents median, minimal and maximal MAE values of the last 10 epochs for each hyperparameter setting. By choosing the model with smallest median MAE in the last 10 epochs we could identify hyperparameter setting, with which the training converged well. Due to GPU space constraints, the maximal batch size was 24 for Model 1 and 9 for Model 4.

For regression Model 1, 2 and 3, training with the MSE loss often diverged for larger learning rate values; this was also the case for Models 2 and 3 with the learning rate values set as proposed in original papers. In general, we observed that training with L1 loss was most stable and produced overall lower MAE values, compared to the use of Mean-Squared Error and Kullback-Leibler divergence losses. Hence, hereafter we used the L1 loss in regression Models 1, 2, 3. The chosen optimal hyperparameter values and the original and resulting model accuracy are given in Supplementary Table 5.

Unless noted otherwise, we used the hyperparameters reported in Supplementary Table 5. in all subsequent experiments. Models based on these hyperparameters represent our baseline models.

### A.3. Execution times

All experiments were run on the same workstation with Intel Core i7-8700K CPU, 64 GB system memory and three NVIDIA GeForce RTX 2080 Ti GPUs, each with 11 GB dedicated memory. The image preprocessing pipelines and model architectures differed based on their execution and training time, respectively, and the hardware requirements (cf. Table 6). The RIG preprocessing pipeline took < 2 minutes, while the more complex AFF+GS took 4-7 minutes per image. The Fs+FSL pipeline was most time consuming, taking > 15 minutes per image on average.

**Table 5:** Proposed hyperparameter values in original literature and the values chosen herein. Only the hyperparameters marked with ^*^ were reevaluated. The resulting model accuracy is reported as MAE in years.

|  | **Model 1** |  | **Model 2** |  |
| --- | --- | --- | --- | --- |
|  | **Proposed** | **Implemented** | **Proposed** | **Implemented** |
| **Input size** | $182 \times 218 \times 182$ | $157 \times 189 \times 170$ | $157 \times 189 \times 15$ | |
| **\*Batch size** | 28 | 16 | 16 | 32 |
| **\*Loss function** | L1 |  | MSE | L1 |
| **\*Learning rate** | $1 \times 10^{-2}$ | $1 \times 10^{-4}$ | $1 \times 10^{-4}$ | $1 \times 10^{-3}$ |
| **Learning rate decay** | 3% | | $1 \times 10^{-4}$ | |
| **Weight decay** | $5 \times 10^{-5}$ | | $1 \times 10^{-3}$ | |
| **Momentum** | 0.9 |  | 0.9 |  |
| **Parameters** | $\approx 900\,000$ | | $\approx 6.6$ mio | |
| **MAE (Test) [years]**<br>*med[$min, max$]* | 4.65 | 3.57 [3.52, 3.61] | 4.0 | 4.23 [4.14, 4.67] |
|  | **Model 3** |  | **Model 4** |  |
|  | **Proposed** | **Implemented** | **Proposed** | **Implemented** |
| **Input size** | $95 \times 79 \times 78$ | | $160 \times 192 \times 160$ | $157 \times 189 \times 170$ |
| **\*Batch size** | 16 | 8 | 8 | 8 |
| **\*Loss function** | MSE | L1 | Kullback-Leibler divergence |  |
| **\*Learning rate** | $5 \times 10^{-5}$ | | $1 \times 10^{-2}$ | |
| **Learning rate decay** | $1 \times 10^{-4}$ | | $\times 0.3$ every 30 epochs | |
| **Weight decay** | $5 \times 10^{-4}$ | | $1 \times 10^{-3}$ | |
| **Momentum** | 0.9 |  | 0.9 |  |
| **Parameters** | $\approx 900\,000$ | | $\approx 6.6$ mio | |
| **MAE (Test) [years]**<br>*med[$min, max$]* | 3.67 | 3.57 [3.52, 4.26] | 2.14 | 3.35 [3.29, 3.42] |

**Table 6:** Average run time of preprocessing pipeline per image (*left*) and model training times with hardware requirements (*right*).

| **Image preprocessing** | **Time [m:ss]** | **Model** | **Time [h]** | **No. GPUs** |
| --- | --- | --- | --- | --- |
| **RIG** | 1:25 | **Model 1** | 15.5 | 2 |
| **RIG+GS** | 6:30 | **Model 2** | 8.9 | 1 |
| **AFF+GS** | 7:40 | **Model 3** | 7.17 | 1 |
| **Fs+FSL** | 16:20 | **Model 4** | 20.2 | 3 |

The difference in both the model training time and the hardware requirements is substantial for different model architectures. Models 1 and 4, trained on full resolution input 3D images require more than twice as much training time and GPU memory, compared to Models 2 and 4. Despite the larger number of trainable parameters in Model 3, its accuracy and robustness were comparable to that of Models 1 and 4.

### A.4. Model predictions on new site

The Supplementary Figure 9 and Supplementary Figure 10 show model predictions on UKB dataset using the same (Section 4.3) and different preprocessing (Section 4.4) as used on the training set. The predictions show a clear systematic offset, specific to each combination of preprocessing and model architecture.

## B. Linear Mixed Effect Model results

The subsequent section presents detailed results from the LMEM and ANOVA tests corresponding to specific experiments. Specifically, refer to Table 7 for Section 4.2, Table 8 for Section 4.3, and Table 9 for Section 4.4. The levels of statistical significance are denoted as: ‘***’ for 0 < *p* < 0.001, ‘**’ for 0.001 < *p* < 0.01 and ‘*’ for 0.01 < *p* < 0.05.

**Table 7:**
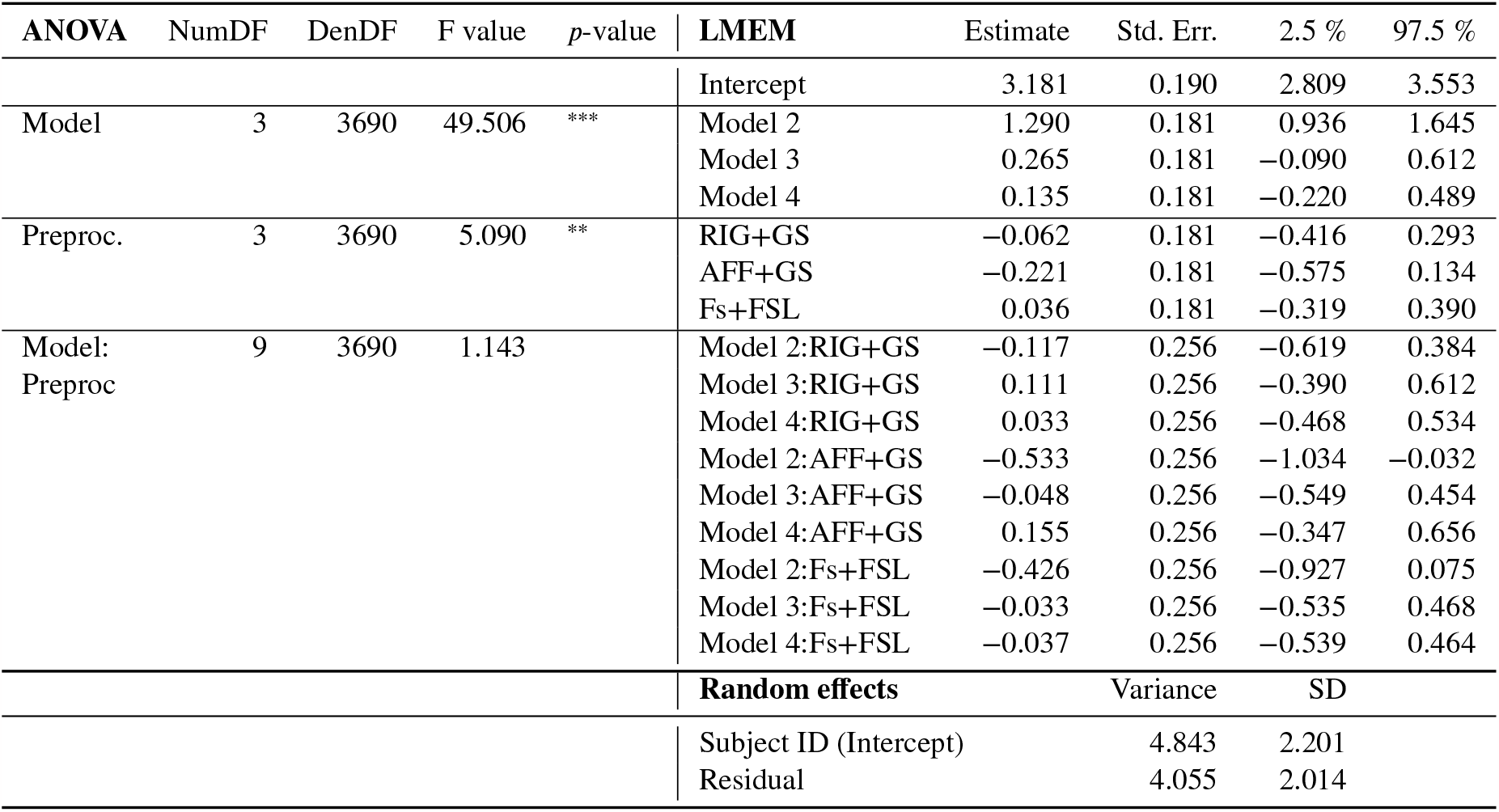
Results of ANOVA and LMEM with absolute error as response variable, and model architecture and preprocessing procedure as fixed factor on test set of Multi-site dataset: *Abs Error* = *Model* + *P reprocessing* + *Model* * *Preprocessing* + (1|*ID*). Interaction was was not statistically significant. Here, ‘NumDF’ denotes the numerator degrees of freedom, and ‘DenDF’ denotes the denominator degrees of freedom.

**Table 8:** Results of ANOVA and LMEM tests on the UK Biobank dataset preprocessed with the same preprocessing procedure as the training dataset with absolute error as response variable, and model architecture, offset correction (OC) and preprocessing procedure as fixed factor: *Abs Error* = *Model* + *P reproc*. + *OC* + *Model* * *P reproc* + *Model* * *OC* + *P reproc* * *OC* + *Model* * *P reproc* * *OC* + (1|*ID*). Here, ‘NumDF’ denotes the numerator degrees of freedom, and ‘DenDF’ denotes the denominator degrees of freedom.

| ANOVA | NumDF | DenDF | F value | p-value | LMEM | Estimate | Std. Err. | 2.5 % | 97.5 % |
| --- | --- | --- | --- | --- | --- | --- | --- | --- | --- |
|  |  |  |  |  | Intercept | 4.478 | 0.082 | 4.317 | 4.640 |
| OC | 1 | 46252 | 637.861 | *** | OC | -1.217 | 0.091 | -1.395 | -1.039 |
| Model | 3 | 46252 | 356.449 | *** | Model 2 | 0.588 | 0.091 | 0.410 | 0.767 |
|  |  |  |  |  | Model 3 | 0.250 | 0.091 | 0.072 | 0.428 |
|  |  |  |  |  | Model 4 | 0.712 | 0.091 | 0.534 | 0.890 |
| Preproc. | 3 | 46252 | 221.364 | *** | RIG+GS | -0.010 | 0.091 | -0.188 | 0.169 |
|  |  |  |  |  | AFF+GS | -0.752 | 0.091 | -0.930 | -0.574 |
|  |  |  |  |  | Fs+FSL | -1.152 | 0.091 | -1.330 | -0.974 |
| OC: | 3 | 46252 | 53.303 | *** | OC:Model 2 | 1.096 | 0.129 | 0.844 | 1.348 |
| Model |  |  |  |  | OC:Model 3 | 0.095 | 0.129 | -0.157 | 0.347 |
|  |  |  |  |  | OC:Model 4 | -0.280 | 0.129 | -0.532 | -0.028 |
| OC: | 3 | 46252 | 93.765 | *** | OC:RIG+GS | 0.077 | 0.129 | -0.175 | 0.329 |
| Preproc. |  |  |  |  | OC:AFF+GS | 0.804 | 0.129 | 0.552 | 1.056 |
|  |  |  |  |  | OC:F<sub>s</sub>+FSL | 1.171 | 0.129 | 0.919 | 1.423 |
| Model: | 9 | 46252 | 12.161 | *** | Model 2:RIG+GS | -0.153 | 0.129 | -0.405 | 0.099 |
| Preproc. |  |  |  |  | Model 3:RIG+GS | 0.543 | 0.129 | 0.291 | 0.795 |
|  |  |  |  |  | Model 4:RIG+GS | -0.687 | 0.129 | -0.939 | -0.435 |
|  |  |  |  |  | Model 2:AFF+GS | 0.010 | 0.129 | -0.242 | 0.262 |
|  |  |  |  |  | Model 3:AFF+GS | -0.047 | 0.129 | -0.299 | 0.205 |
|  |  |  |  |  | Model 4:AFF+GS | -0.789 | 0.129 | -1.041 | -0.537 |
|  |  |  |  |  | Model 2:F<sub>s</sub>+FSL | 0.628 | 0.129 | 0.376 | 0.880 |
|  |  |  |  |  | Model 3:F<sub>s</sub>+FSL | 0.207 | 0.129 | -0.045 | 0.459 |
|  |  |  |  |  | Model 4:F<sub>s</sub>+FSL | -0.410 | 0.129 | -0.662 | -0.158 |
| OC: | 9 | 46252 | 12.981 | *** | OC:Model 2:RIG+GS | -0.113 | 0.182 | -0.470 | 0.243 |
| Model: |  |  |  |  | OC:Model 3:RIG+GS | -0.513 | 0.182 | -0.869 | -0.156 |
| Preproc. |  |  |  |  | OC:Model 4:RIG+GS | 0.634 | 0.182 | 0.277 | 0.990 |
|  |  |  |  |  | OC:Model 2:AFF+GS | -0.796 | 0.182 | -1.152 | -0.439 |
|  |  |  |  |  | OC:Model 3:AFF+GS | -0.096 | 0.182 | -0.453 | 0.260 |
|  |  |  |  |  | OC:Model 4:AFF+GS | 0.491 | 0.182 | 0.134 | 0.847 |
|  |  |  |  |  | OC:Model 2:F<sub>s</sub>+FSL | -1.052 | 0.182 | -1.408 | -0.695 |
|  |  |  |  |  | OC:Model 3:F<sub>s</sub>+FSL | -0.410 | 0.182 | -0.766 | -0.053 |
|  |  |  |  |  | OC:Model 4:F<sub>s</sub>+FSL | 0.285 | 0.182 | -0.072 | 0.641 |
|  |  |  |  |  | **Random effects** | Variance |  | SD |  |
|  |  |  |  |  | Subject ID (Intercept) | 3.937 |  | 1.984 |  |
|  |  |  |  |  | Residual | 6.174 |  | 2.485 |  |

**Table 9:** Results of ANOVA and LMEM tests on UK Biobank dataset preprocessed with new preprocessing procedure with absolute error as response variable, and model architecture, offset correction (OC) and preprocessing procedure as fixed factor: *Abs Error* = *OC* + *Model* + *Preprocessing* + *OC* * *Model* + *OC* * *P reprocessing* + *Model* * *Preprocessing* +*Model* * *Preprocessing* * *OC* +(1|*ID*). Here, ‘NumDF’ denotes the numerator degrees of freedom, and ‘DenDF’ denotes the denominator degrees of freedom.

| ANOVA | NumDF | DenDF | F value | p-value | LMEM | Estimate | Std. Err. | 2.5 % | 97.5 % |
| --- | --- | --- | --- | --- | --- | --- | --- | --- | --- |
|  |  |  |  |  | Intercept | 9.228 | 0.107 | 9.019 | 9.436 |
| OC | 1 | 46252 | 20035.70 | *** | OC | -5.588 | 0.129 | -5.840 | -5.335 |
| Model | 3 | 46252 | 2514.49 | *** | Model 2 | 3.681 | 0.129 | 3.428 | 3.933 |
|  |  |  |  |  | Model 3 | -0.694 | 0.129 | -0.947 | -0.442 |
|  |  |  |  |  | Model 4 | -1.852 | 0.129 | -2.104 | -1.599 |
| Preproc | 3 | 46252 | 3513.01 | *** | RIG+GS | 1.295 | 0.129 | 1.043 | 1.548 |
|  |  |  |  |  | AFF+GS | -4.574 | 0.129 | -4.826 | -4.321 |
|  |  |  |  |  | Fs+FSL | -5.003 | 0.129 | -5.256 | -4.751 |
| OC: | 3 | 46252 | 946.12 | *** | OC:Model 2 | -1.737 | 0.182 | -2.094 | -1.380 |
| Model |  |  |  |  | OC:Model 3 | 0.648 | 0.182 | 0.291 | 1.005 |
|  |  |  |  |  | OC:Model 4 | 2.513 | 0.182 | 2.156 | 2.870 |
| OC: | 3 | 46252 | 3126.95 | *** | OC:RIG+GS | -1.227 | 0.182 | -1.585 | -0.870 |
| Preproc |  |  |  |  | OC:AFF+GS | 4.648 | 0.182 | 4.291 | 5.005 |
|  |  |  |  |  | OC:F<sub>s</sub>+FSL | 5.027 | 0.182 | 4.670 | 5.384 |
| Model: | 9 | 46252 | 77.89 | *** | Model 2:RIG+GS | 3.996 | 0.182 | 3.639 | 4.354 |
| Preproc |  |  |  |  | Model 3:RIG+GS | 4.631 | 0.182 | 4.274 | 4.988 |
|  |  |  |  |  | Model 4:RIG+GS | 3.134 | 0.182 | 2.777 | 3.491 |
|  |  |  |  |  | Model 2:AFF+GS | 1.466 | 0.182 | 1.109 | 1.823 |
|  |  |  |  |  | Model 3:AFF+GS | 3.978 | 0.182 | 3.621 | 4.335 |
|  |  |  |  |  | Model 4:AFF+GS | 1.632 | 0.182 | 1.275 | 1.989 |
|  |  |  |  |  | Model 2:F<sub>s</sub>+FSL | 0.979 | 0.182 | 0.622 | 1.336 |
|  |  |  |  |  | Model 3:F<sub>s</sub>+FSL | 0.920 | 0.182 | 0.563 | 1.277 |
|  |  |  |  |  | Model 4:F<sub>s</sub>+FSL | 1.456 | 0.182 | 1.099 | 1.813 |
| OC: | 9 | 46252 | 71.25 | *** | OC:Model 2:RIG+GS | -4.394 | 0.258 | -4.899 | -3.889 |
| Model: |  |  |  |  | OC:Model 3:RIG+GS | -4.410 | 0.258 | -4.915 | -3.905 |
| Preproc |  |  |  |  | OC:Model 4:RIG+GS | -2.594 | 0.258 | -3.099 | -2.090 |
|  |  |  |  |  | OC:Model 2:AFF+GS | -2.443 | 0.258 | -2.948 | -1.938 |
|  |  |  |  |  | OC:Model 3:AFF+GS | -3.835 | 0.258 | -4.340 | -3.330 |
|  |  |  |  |  | OC:Model 4:AFF+GS | -1.933 | 0.258 | -2.438 | -1.428 |
|  |  |  |  |  | OC:Model 2:F<sub>s</sub>+FSL | -0.686 | 0.258 | -1.191 | -0.181 |
|  |  |  |  |  | OC:Model 3:F<sub>s</sub>+FSL | -0.787 | 0.258 | -1.292 | -0.282 |
|  |  |  |  |  | OC:Model 4:F<sub>s</sub>+FSL | -1.951 | 0.258 | -2.456 | -1.446 |
|  |  |  |  |  | **Random effects** | Variance |  | SD |  |
|  |  |  |  |  | Subject ID (Intercept) | 4.577 |  | 2.139 |  |
|  |  |  |  |  | Residual | 12.396 |  | 3.521 |  |

## Declaration of generative AI and AI-assisted technologies in the writing process

During the preparation of this work the author(s) used ChatGPT-4 in order to improve the readability of this paper. After using this tool/service, the author(s) reviewed and edited the content as needed and take(s) full responsibility for the content of the publication.

## CRediT authorship contribution statement

**Lara Dular:** Conceptualization and evaluation protocol of this study, Data cleanup, Implementation of methods and experiments, Collection and analysis of results, Wrote and revised the manuscript. **Franjo Pernuš:** Wrote and revised the manuscript. **Žiga Špiclin:** Conceptualization and evaluation protocol of this study, Data collection, Wrote and revised the manuscript.

## Footnotes

1 Adaptive non-local means denoising implementation: https://github.com/djkwon/naonlm3d

2 N4 bias field correction: https://manpages.debian.org/testing/ants/N4BiasFieldCorrection.1.en.html

3 NiftyReg Software http://cmictig.cs.ucl.ac.uk/wiki/index.php/NiftyReg

4 Freesurfer: https://surfer.nmr.mgh.harvard.edu/

5 FSL (FMRIB Software Library): https://fsl.fmrib.ox.ac.uk/fsl/fslwiki

6 The application of weighted training led to a statistically significant reduction in absolute error for subjects over the age of 80 years, where the number of training samples is lower. This significance was exclusively observed for Model 4 (*p* < 0.001), whereas other models didn’t show such an effect (results not shown).

7 Some public data sources may require online registration to gain access to the T1w MRI scans. The UKB dataset is available for a fee.

## References

K. Franke, C. Gaser, Ten Years of BrainAGE as a Neuroimaging Biomarker of Brain Aging: What Insights Have We Gained?, Front Neurol 10 (2019) 789.

K. Ning, L. Zhao, W. Matloff, F. Sun, A. W. Toga, Association of relative brain age with tobacco smoking, alcohol consumption, and genetic variants, Sci Rep 10 (2020) 10.

Z. Linli, J. Feng, W. Zhao, S. Guo, Associations between smoking and accelerated brain ageing, Progress in Neuro-Psychopharmacology and Biological Psychiatry 113 (2022) 110471.

J. H. Cole, Multimodality neuroimaging brain-age in UK biobank: relationship to biomedical, lifestyle, and cognitive factors, Neurobiology of Aging 92 (2020) 34–42.

K. Franke, C. Gaser, B. Manor, V. Novak, Advanced BrainAGE in older adults with type 2 diabetes mellitus, Front Aging Neurosci 5 (2013).

N. Bittner, C. Jockwitz, K. Franke, C. Gaser, S. Moebus, U. J. Bayen, K. Amunts, S. Caspers, When your brain looks older than expected: combined lifestyle risk and BrainAGE, Brain Struct Funct 226 (2021) 621–645.

S. M. Smith, D. Vidaurre, F. Alfaro-Almagro, T. E. Nichols, K. L. Miller, Estimation of brain age delta from brain imaging, NeuroImage 200 (2019) 528–539.

M. Habes, R. Pomponio, H. Shou, J. Doshi, E. Mamourian, G. Erus, I. Nasrallah, L. J. Launer, T. Rashid, M. Bilgel, Y. Fan, J. B. Toledo, K. Yaffe, A. Sotiras, D. Srinivasan, M. Espeland, C. Masters, P. Maruff, J. Fripp, H. Völzk, S. C. Johnson, J. C. Morris, M. S. Albert, M. I. Miller, R. N. Bryan, H. J. Grabe, S. M. Resnick, D. A. Wolk, C. Davatzikos, for the iSTAGING consortium, the Preclinical AD consortium, the ADNI, and the CARDIA studies, The Brain Chart of Aging: Machine-learning analytics reveals links between brain aging, white matter disease, amyloid burden, and cognition in the iSTAGING consortium of 10,216 harmonized MR scans, Alzheimer’s & Dementia 17 (2021) 89–102.

P. Jawinski, S. Markett, J. Drewelies, S. Düzel, I. Demuth, E. Steinhagen-Thiessen, G. G. Wagner, D. Gerstorf, U. Lindenberger, C. Gaser, S. Kühn, Linking brain age gap to mental and physical health in the berlin aging study ii, Frontiers in Aging Neuroscience 14 (2022).

F. Liem, G. Varoquaux, J. Kynast, F. Beyer, S. Kharabian Masouleh, J. M. Huntenburg, L. Lampe, M. Rahim, A. Abraham, R. C. Craddock, S. Riedel-Heller, T. Luck, M. Loeffler, M. L. Schroeter, A. V. Witte, A. Villringer, D. S. Margulies, Predicting brain-age from multimodal imaging data captures cognitive impairment, NeuroImage 148 (2017) 179–188.

R. Lay-Yee, A. R. Hariri, A. R. Knodt, A. Barrett-Young, T. Matthews, B. J. Milne, Social isolation from childhood to mid-adulthood: is there an association with older brain age?, Psychological Medicine (2023) 1–9.

K. Franke, C. Gaser, Longitudinal Changes in Individual BrainAGE in Healthy Aging, Mild Cognitive Impairment, and Alzheimer’s Disease, GeroPsych 25 (2012) 235–245.

E. A. Høgestøl, T. Kaufmann, G. O. Nygaard, M. K. Beyer, P. Sowa, J. E. Nordvik, K. Kolskår, G. Richard, O. A. Andreassen, H. F. Harbo, L. T. Westlye, Cross-Sectional and Longitudinal MRI Brain Scans Reveal Accelerated Brain Aging in Multiple Sclerosis, Frontiers in Neurology 10 (2019).

J. H. Cole, J. Raffel, T. Friede, A. Eshaghi, W. J. Brownlee, D. Chard, N. D. Stefano, C. Enzinger, L. Pirpamer, M. Filippi, C. Gasperini, M. A. Rocca, A. Rovira, S. Ruggieri, J. Sastre-Garriga, M. L. Stromillo, B. M. J. Uitdehaag, H. Vrenken, F. Barkhof, R. Nicholas, O. Ciccarelli, Longitudinal Assessment of Multiple Sclerosis with the Brain-Age Paradigm, Annals of Neurology 88 (2020) 93–105.

H. G. Schnack, N. E. van Haren, M. Nieuwenhuis, H. E. Hulshoff Pol, W. Cahn, R. S. Kahn, Accelerated Brain Aging in Schizophrenia: A Longitudinal Pattern Recognition Study, AJP 173 (2016) 607–616.

N. Koutsouleris, C. Davatzikos, S. Borgwardt, C. Gaser, R. Bottlender, T. Frodl, P. Falkai, A. Riecher-Rössler, H.-J. Möller, M. Reiser, C. Pantelis, E. Meisenzahl, Accelerated brain aging in schizophrenia and beyond: a neuroanatomical marker of psychiatric disorders, Schizophr Bull 40 (2014) 1140–1153.

K. J. Petersen, N. Metcalf, S. Cooley, D. Tomov, F. Vaida, R. Paul, B. M. Ances, Accelerated Brain Aging and Cerebral Blood Flow Reduction in Persons With Human Immunodeficiency Virus, Clinical Infectious Diseases 73 (2021) 1813–1821.

J. H. Cole, J. Underwood, M. W. A. Caan, D. D. Francesco, R. A. v. Zoest, R. Leech, F. W. N. M. Wit, P. Portegies, G. J. Geurtsen, B. A. Schmand, M. F. S. v. d. Loeff, C. Franceschi, C. A. Sabin, C. B. L. M. Majoie, A. Winston, P. Reiss, D. J. Sharp, Increased brain-predicted aging in treated HIV disease, Neurology 88 (2017) 1349–1357.

D. M. Hedderich, A. Menegaux, B. Schmitz-Koep, R. Nuttall, J. Zimmermann, S. C. Schneider, J. G. Bäuml, M. Daamen, H. Boecker, M. Wilke, C. Zimmer, D. Wolke, P. Bartmann, C. Sorg, C. Gaser, Increased Brain Age Gap Estimate (BrainAGE) in Young Adults After Premature Birth, Front. Aging Neurosci. 13 (2021).

G. Richard, K. Kolskår, A.-M. Sanders, T. Kaufmann, A. Petersen, N. T. Doan, J. M. Sánchez, D. Alnæs, K. M. Ulrichsen, E. S. Dørum, O. A. Andreassen, J. E. Nordvik, L. T. Westlye, Assessing distinct patterns of cognitive aging using tissue-specific brain age prediction based on diffusion tensor imaging and brain morphometry, PeerJ 6 (2018) e5908.

I. Hwang, E. K. Yeon, J. Y. Lee, R.-E. Yoo, K. M. Kang, T. J. Yun, S. H. Choi, C.-H. Sohn, H. Kim, J.-h. Kim, Prediction of brain age from routine T2-weighted spin-echo brain magnetic resonance images with a deep convolutional neural network, Neurobiology of Aging 105 (2021) 78–85.

J. Gao, J. Liu, Y. Xu, D. Peng, Z. Wang, Brain age prediction using the graph neural network based on resting-state functional MRI in Alzheimer’s disease, Front Neurosci 17 (2023) 1222751.

L. Baecker, R. Garcia-Dias, S. Vieira, C. Scarpazza, A. Mechelli, Machine learning for brain age prediction: Introduction to methods and clinical applications, eBioMedicine 72 (2021).

P. K. Lam, V. Santhalingam, P. Suresh, R. Baboota, A. H. Zhu, S. I. Thomopoulos, N. Jahanshad, P. M. Thompson, Accurate brain age prediction using recurrent slice-based networks, in: J. Brieva, N. Lepore, M. G. Linguraru, E. R. C. M.D. (Eds.), 16th International Symposium on Medical Information Processing and Analysis, volume 11583, International Society for Optics and Photonics, SPIE, 2020, p. 1158303. URL: 10.1117/12.2579630. doi:10.1117/12.2579630.

H. Peng, W. Gong, C. F. Beckmann, A. Vedaldi, S. M. Smith, Accurate brain age prediction with lightweight deep neural networks, Medical Image Analysis 68 (2021).

B. Dufumier, P. Gori, I. Battaglia, J. Victor, A. Grigis, E. Duchesnay, Benchmarking cnn on 3d anatomical brain mri: Architectures, data augmentation and deep ensemble learning, 2021. URL: https://arxiv.org/abs/2106.01132. doi:10.48550/ARXIV.2106.01132.

X. Feng, Z. C. Lipton, J. Yang, S. A. Small, F. A. Provenzano, Estimating brain age based on a uniform healthy population with deep learning and structural magnetic resonance imaging, Neurobiology of Aging 91 (2020) 15–25.

C. Dartora, A. Marseglia, G. Mårtensson, G. Rukh, J. Dang, J.-S. Muehlboeck, L.-O. Wahlund, R. Moreno, J. Barroso, D. Ferreira, H. B. Schiöth, E. Westman, A. D. N. Initiative, A. I. Biomarkers, L. flagship study of ageing, J. A. D. N. Initiative, A. consortium, A deep learning model for brain age prediction using minimally pre-processed t1w-images as input, medRxiv (2023).

J. H. Cole, R. P. K. Poudel, D. Tsagkrasoulis, M. W. A. Caan, C. Steves, T. D. Spector, G. Montana, Predicting brain age with deep learning from raw imaging data results in a reliable and heritable biomarker, NeuroImage 163 (2017) 115–124.

M. Ueda, K. Ito, K. Wu, K. Sato, Y. Taki, H. Fukuda, T. Aoki, An Age Estimation Method Using 3D-CNN From Brain MRI Images, in: 2019 IEEE 16th International Symposium on Biomedical Imaging (ISBI 2019), 2019, pp. 380–383. doi:10.1109/ISBI.2019.8759392.

T.-W. Huang, H.-T. Chen, R. Fujimoto, K. Ito, K. Wu, K. Sato, Y. Taki, H. Fukuda, T. Aoki, Age estimation from brain MRI images using deep learning, in: 2017 IEEE 14th International Symposium on Biomedical Imaging (ISBI 2017), 2017, pp. 849–852. doi:10.1109/ISBI.2017.7950650.

K.-M. Bintsi, V. Baltatzis, A. Kolbeinsson, A. Hammers, D. Rueckert, Patch-based Brain Age Estimation from MR Images, 2020. URL: http://arxiv.org/abs/2008.12965.

J. Cheng, Z. Liu, H. Guan, Z. Wu, H. Zhu, J. Jiang, W. Wen, D. Tao, T. Liu, Brain age estimation from mri using cascade networks with ranking loss, IEEE Transactions on Medical Imaging 40 (2021) 3400–3412.

L. Fisch, J. Ernsting, N. R. Winter, V. Holstein, R. Leenings, M. Beisemann, K. Sarink, D. Emden, N. Opel, R. Redlich, J. Repple, D. Grotegerd, S. Meinert, N. Wulms, H. Minnerup, J. G. Hirsch, T. Niendorf, B. Endemann, F. Bamberg, T. Kröncke, A. Peters, R. Bülow, H. Völzke, O. von Stackelberg, R. F. Sowade, L. Umutlu, B. Schmidt, S. Caspers, Consortium, German National Cohort Study Center, H. Kugel, B. T. Baune, T. Kircher, B. Risse, U. Dannlowski, K. Berger, T. Hahn, Predicting brain-age from raw t 1 -weighted magnetic resonance imaging data using 3d convolutional neural networks, 2021. URL: https://arxiv.org/abs/2103.11695. doi:10.48550/ARXIV.2103.11695.

S. Lathuilière, P. Mesejo, X. Alameda-Pineda, R. Horaud, A comprehensive analysis of deep regression, IEEE Trans. Pattern Anal. Mach. Intell. 42 (2020) 2065–2081.

S. Kharabian Masouleh, S. B. Eickhoff, Y. Zeighami, L. B. Lewis, R. Dahnke, C. Gaser, F. Chouinard-Decorte, C. Lepage, L. H. Scholtens, F. Hoffstaedter, D. C. Glahn, J. Blangero, A. C. Evans, S. Genon, S. L. Valk, Influence of Processing Pipeline on Cortical Thickness Measurement, Cereb Cortex 30 (2020) 5014–5027.

N. Bhagwat, A. Barry, E. W. Dickie, S. T. Brown, G. A. Devenyi, K. Hatano, E. DuPre, A. Dagher, M. Chakravarty, C. M. T. Greenwood, B. Misic, D. N. Kennedy, J.-B. Poline, Understanding the impact of preprocessing pipelines on neuroimaging cortical surface analyses, GigaScience 10 (2021).

M. de Fátima Machado Dias, P. Carvalho, M. Castelo-Branco, J. Valente Duarte, Cortical thickness in brain imaging studies using freesurfer and cat12: A matter of reproducibility, Neuroimage: Reports 2 (2022) 100137.

M. Tanveer, M. Ganaie, I. Beheshti, T. Goel, N. Ahmad, K.-T. Lai, K. Huang, Y.-D. Zhang, J. Del Ser, C.-T. Lin, Deep learning for brain age estimation: A systematic review, Information Fusion 96 (2023) 130–143.

B. A. Jonsson, G. Bjornsdottir, T. E. Thorgeirsson, L. M. Ellingsen, G. B. Walters, D. F. Gudbjartsson, H. Stefansson, K. Stefansson, M. O. Ulfarsson, Brain age prediction using deep learning uncovers associated sequence variants, Nat Commun 10 (2019) 5409.

J. V. Manjón, P. Coupé, L. Martí-Bonmatí, D. L. Collins, M. Robles, Adaptive non-local means denoising of MR images with spatially varying noise levels, J Magn Reson Imaging 31 (2010) 192–203.

V. Fonov, A. Evans, R. McKinstry, C. Almli, D. Collins, Unbiased nonlinear average age-appropriate brain templates from birth to adulthood, NeuroImage 47 (2009) S102.

N. J. Tustison, B. B. Avants, P. A. Cook, Y. Zheng, A. Egan, P. A. Yushkevich, J. C. Gee, N4ITK: improved N3 bias correction, IEEE Trans Med Imaging 29 (2010) 1310–1320.

M. Modat, D. M. Cash, P. Daga, G. P. Winston, J. S. Duncan, S. Ourselin, Global image registration using a symmetric block-matching approach, Journal of Medical Imaging 1 (2014) 1 –6.

M. Jenkinson, C. F. Beckmann, T. E. J. Behrens, M. W. Woolrich, S. M. Smith, FSL, NeuroImage 62 (2012) 782–790.

Laboratory for Computational Neuroimaging, Athinoula A. Martinos Center for Biomedical Imaging., Freesurferwiki: recon-all, 2022. URL: https://surfer.nmr.mgh.harvard.edu/fswiki/recon-all.

M. Jenkinson, P. Bannister, M. Brady, S. Smith, Improved optimization for the robust and accurate linear registration and motion correction of brain images, Neuroimage 17 (2002) 825–841.

S. M. Smith, F. Alfaro-Almagro, K. L. Miller, UK Biobank Brain Imaging Documentation, Welcome Centre for Integrative Neuroimaging and Oxford University, 2020. URL: https://biobank.ctsu.ox.ac.uk/crystal/crystal/docs/brain_mri.pdf.

G. Grabner, A. L. Janke, M. M. Budge, D. Smith, J. Pruessner, D. L. Collins, Symmetric Atlasing and Model Based Segmentation: An Application to the Hippocampus in Older Adults, in: R. Larsen, M. Nielsen, J. Sporring (Eds.), Medical Image Computing and Computer-Assisted Intervention – MICCAI 2006, Springer Berlin Heidelberg, Berlin, Heidelberg, 2006, pp. 58–66.

M. Jenkinson, S. Smith, A global optimisation method for robust affine registration of brain images, Med Image Anal 5 (2001) 143–156.

A.-M. G. d. Lange, T. Kaufmann, D. v. d. Meer, L. A. Maglanoc, D. Alnæs, T. Moberget, G. Douaud, O. A. Andreassen, L. T. Westlye, Population-based neuroimaging reveals traces of childbirth in the maternal brain, PNAS 116 (2019) 22341–22346.

J. H. Cole, T. Annus, L. R. Wilson, R. Remtulla, Y. T. Hong, T. D. Fryer, J. Acosta-Cabronero, A. Cardenas-Blanco, R. Smith, D. K. Menon, S. H. Zaman, P. J. Nestor, A. J. Holland, Brain-predicted age in Down syndrome is associated with beta amyloid deposition and cognitive decline, Neurobiology of Aging 56 (2017) 41–49.

T. Dunås, A. Wåhlin, L. Nyberg, C.-J. Boraxbekk, Multimodal Image Analysis of Apparent Brain Age Identifies Physical Fitness as Predictor of Brain Maintenance, Cerebral Cortex (2021).

E. R. Butler, A. Chen, R. Ramadan, T. T. Le, K. Ruparel, T. M. Moore, T. D. Satterthwaite, F. Zhang, H. Shou, R. C. Gur, T. E. Nichols, R. T. Shinohara, Pitfalls in brain age analyses, Human Brain Mapping 42 (2021) 4092–4101.

A.-M. G. de Lange, M. Anatürk, J. Rokicki, L. K. M. Han, K. Franke, D. Alnæs, K. P. Ebmeier, B. Draganski, T. Kaufmann, L. T. Westlye, T. Hahn, J. H. Cole, Mind the gap: Performance metric evaluation in brain-age prediction, Human Brain Mapping 43 (2022) 3113–3129.

G. Levakov, G. Rosenthal, I. Shelef, T. R. Raviv, G. Avidan, From a deep learning model back to the brain—Identifying regional predictors and their relation to aging, Human Brain Mapping 41 (2020) 3235–3252.

C.-Y. Kuo, T.-M. Tai, P.-L. Lee, C.-W. Tseng, C.-Y. Chen, L.-K. Chen, C.-K. Lee, K.-H. Chou, S. See, C.-P. Lin, Improving Individual Brain Age Prediction Using an Ensemble Deep Learning Framework, Front Psychiatry 12 (2021).

J. Jönemo, M. U. Akbar, R. Kämpe, J. P. Hamilton, A. Eklund, Efficient brain age prediction from 3d mri volumes using 2d projections, 2022. URL: https://arxiv.org/abs/2211.05762. doi:10.48550/ARXIV.2211.05762.

J. H. Cole, K. Franke, N. Cherbuin, Quantification of the Biological Age of the Brain Using Neuroimaging, in: A. Moskalev (Ed.), Biomarkers of Human Aging, Healthy Ageing and Longevity, Springer International Publishing, Cham, 2019, pp. 293–328. URL: 10.1007/978-3-030-24970-0_19. doi:10.1007/978-3-030-24970-0_19.

M. A. Shafto, L. K. Tyler, M. Dixon, J. R. Taylor, J. B. Rowe, R. Cusack, A. J. Calder, W. D. Marslen-Wilson, J. Duncan, T. Dalgleish, R. N. Henson, C. Brayne, F. E. Matthews, The Cambridge Centre for Ageing and Neuroscience (Cam-CAN) study protocol: a cross-sectional, lifespan, multidisciplinary examination of healthy cognitive ageing, BMC Neurol 14 (2014).

J. R. Taylor, N. Williams, R. Cusack, T. Auer, M. A. Shafto, M. Dixon, L. K. Tyler, n. Cam-Can, R. N. Henson, The Cambridge Centre for Ageing and Neuroscience (Cam-CAN) data repository: Structural and functional MRI, MEG, and cognitive data from a cross-sectional adult lifespan sample, Neuroimage 144 (2017) 262–269.

R. Souza, O. Lucena, J. Garrafa, D. Gobbi, M. Saluzzi, S. Appenzeller, L. Rittner, R. Frayne, R. Lotufo, An open, multi-vendor, multi-field-strength brain MR dataset and analysis of publicly available skull stripping methods agreement, NeuroImage 170 (2018) 482 –494.

D. S. Marcus, A. F. Fotenos, J. G. Csernansky, J. C. Morris, R. L. Buckner, Open Access Series of Imaging Studies: Longitudinal MRI Data in Nondemented and Demented Older Adults, Journal of Cognitive Neuroscience 22 (2010) 2677–2684.

K. L. Miller, F. Alfaro-Almagro, N. K. Bangerter, D. L. Thomas, E. Yacoub, J. Xu, A. J. Bartsch, S. Jbabdi, S. N. Sotiropoulos, J. L. R. Andersson, L. Griffanti, G. Douaud, T. W. Okell, P. Weale, I. Dragonu, S. Garratt, S. Hudson, R. Collins, M. Jenkinson, P. M. Matthews, S. M. Smith, Multimodal population brain imaging in the UK Biobank prospective epidemiological study, Nat Neurosci 19 (2016) 1523–1536.

